# Conformational dynamics in specialized C_2_H_2_ zinc finger domains enable zinc-responsive gene repression in S. pombe

**DOI:** 10.1101/2024.09.20.614115

**Authors:** Vibhuti Wadhwa, Cameron Jamshidi, Kye Stachowski, Amanda J. Bird, Mark P. Foster

## Abstract

Loz1 is a zinc-responsive transcription factor in fission yeast that maintains cellular zinc homeostasis by repressing the expression of genes required for zinc uptake in high zinc conditions. Previous deletion analysis of Loz1 found a region containing two tandem C_2_H_2_ zinc-fingers and an upstream “accessory domain” rich in histidine, lysine, and arginine residues to be sufficient for zinc-dependent DNA binding and gene repression. Here we report unexpected biophysical properties of this pair of seemingly classical C_2_H_2_ zinc fingers. Isothermal titration calorimetry and NMR spectroscopy reveal two distinct zinc binding events localized to the zinc fingers. NMR spectra reveal complex dynamic behavior in this zinc responsive region spanning time scales from fast 10^−12^-10^−10^ to slow > 10^0^ sec. Slow exchange due to *cis-trans* isomerization of the TGERP linker results in doubling of many signals in the protein.

Conformational exchange on the 10^−3^ s timescale throughout the first zinc finger distinguishes it from the second and is linked to weaker affinity for zinc. These findings reveal the mechanism of zinc sensing by Loz1 and illuminate how the protein’s rough free-energy landscape enable zinc sensing, DNA binding and regulated gene expression.

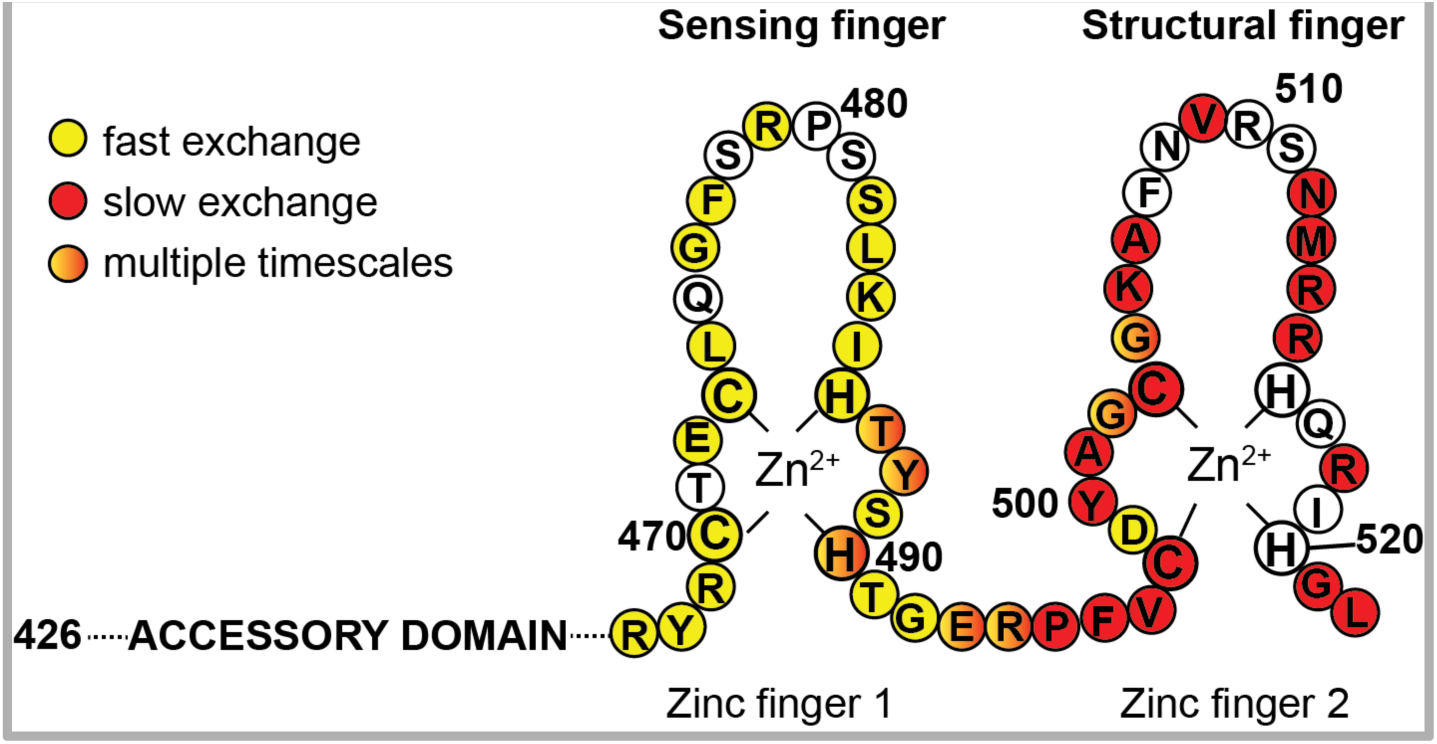

## Introduction

Zinc is an essential nutrient for all living organisms and is the second most abundant trace metal in the human body.^1^ Zinc plays a catalytic role as a Lewis acid in many enzymes and confers structural stability in a variety of proteins. One of the most prevalent structural domains in eukaryotic transcriptional factors is the “zinc finger” (ZF)^2^. Classical C_2_H_2_-type ZFs are ∼30-residue domains which form a ββα fold stabilized by a single zinc ion that is coordinated through side chain atoms of conserved cysteine and histidine residues. C_2_H_2_ ZFs have a consensus sequence of φ-X-Cys-X(2-5)-Cys-X(3)-φ-X(5)-φ-X(2)-His-X(2-5)-H, where φ is hydrophobic amino acid and X is any amino acid. The conserved hydrophobic residues stabilize the core of the each individual ZF domain. In most C_2_H_2_ ZF proteins, two or more ZFs are arranged in tandem through a canonical TGE(R/K)P linker^3,4^

Because canonical C_2_H_2_ ZF domains typically form very stable scaffolds and have high zinc binding affinity (equilibrium dissociation constants K_d_ ranging from 10^−9^ to 10^−12^ M),^5^ zinc binding by these domains is generally associated with a structural role.^6^ However, in a few transcription factors that help cells to maintain zinc homeostasis ZF domains have been found to function as sensors of intracellular zinc levels ^7–9^. In mammals, fish, and reptiles the metal-responsive transcription factor (MTF-1) activates target gene expression when zinc is in excess^10^. The six C_2_H_2_-type ZFs, part of the DNA-binding domain of MTF-1, show increased DNA binding activity upon zinc supplementation^10,11^. Experimental evidence suggests that differential zinc-binding affinity by the six ZF domains underlies zinc-sensing and gene regulation by MTF-1. However, structure-function studies to identify the metalloregulatory finger/subset of fingers of MTF-1 have yielded conflicting observations^12–14^.

Another well characterized zinc-responsive transcription factor is the yeast *Saccharomyces cerevisiae* protein Zap1, which upregulates expression of genes involved in zinc uptake under zinc-limiting conditions. Zap1 is regulated at both the transcriptional and post-translational level by zinc, and contains multiple zinc-regulatory domains which can function independently of each other.^15,16^ The most widely studied transcriptional activation domain (AD2) contains two non-canonical, CWCH_2_-type zinc fingers. Upon zinc binding and folding, the zinc finger pair interact through a network of inter-finger hydrophobic contacts to form a single structural unit, which is hypothesized to be responsible for reduced AD2 activity and transcription.^8,17,18^

In *Schizosaccharomyces pombe* (*Spo*), the transcription factor Loz1 (Loss of Zinc sensing 1) plays a central role in zinc homeostasis ^19^. Under conditions of excess zinc, Loz1 is recruited to the Loz1 response promoter element (LRE, 5’-CGNMGATCNTY-3’; N, any base; M, A or C; Y, pyrimidine), which is both required and sufficient for Loz1-mediated gene repression^20^. Loz1 represses expression of ∼ 30 genes, including *zrt1*, which encodes a high affinity zinc uptake transporter, and *adh1AS*, an antisense transcript that controls the expression of the zinc-dependent alcohol dehydrogenase gene *adh1*^20^. Loz1 also negatively regulates its own expression.^20,21^ An N-terminal deletion construct of this protein (residues 426-522, here termed Loz1AZZ; Figure 1), containing a pair of classical C_2_H_2_ ZFs at the extreme C-terminus, and an upstream 40-amino acid accessory domain is sufficient for DNA binding *in vitro* and can confer partial zinc-dependent repression *in vivo.*^21,22^ Mutations in the ZF domains that disrupt DNA binding abolish zinc-dependent activity of Loz1.^22^

**Figure 1.**
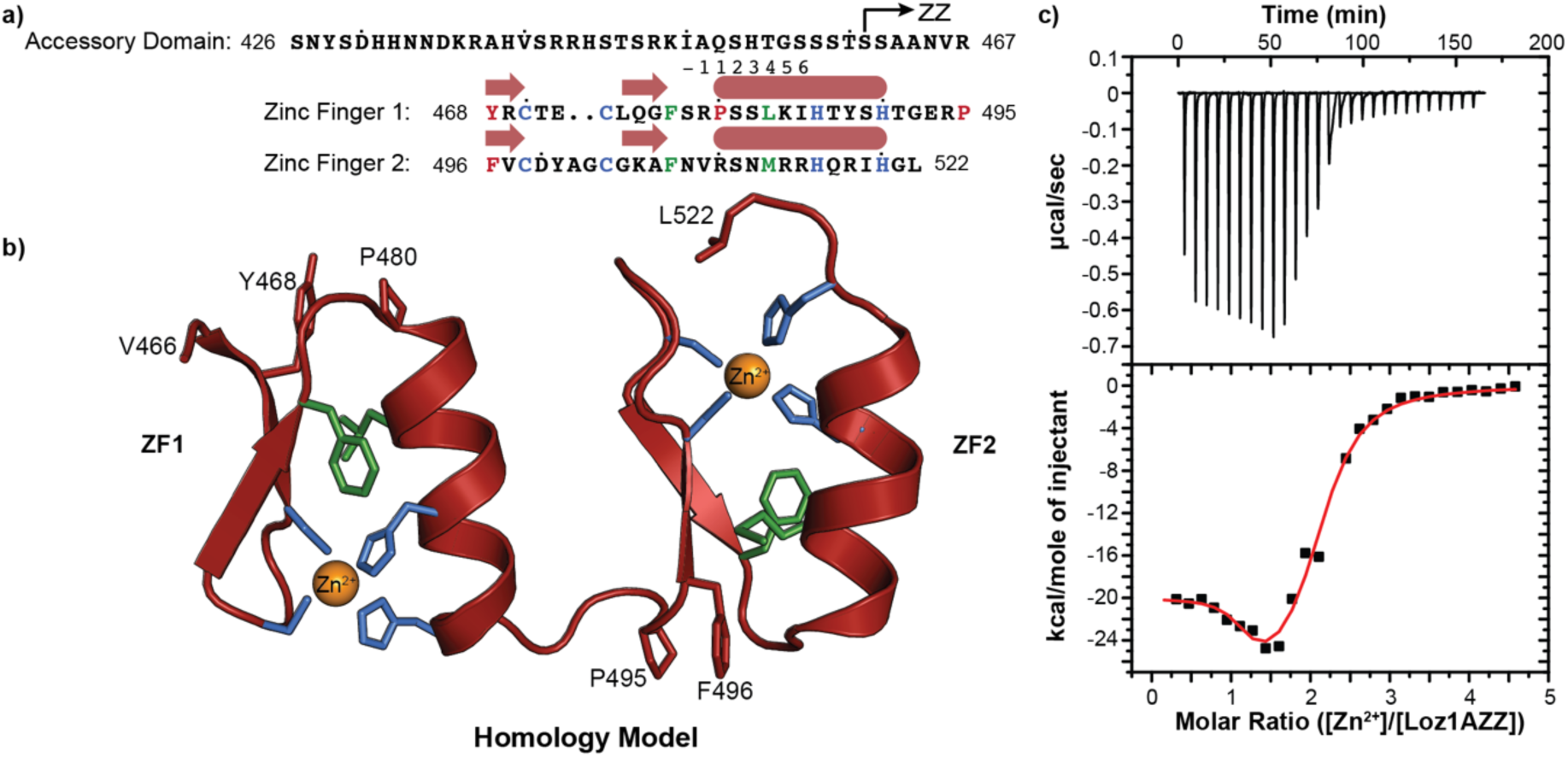
The Loz1 zinc-responsive element binds two zinc ions with different affinities. **a)** The amino acid sequence of Loz1AZZ (residues 426-522) with Zn^2+^-coordinating residues blue and conserved hydrophobic core residues green. The black arrow marks the first residue of the Loz1ZZ construct lacking most of the accessory domain. The secondary structure schematic represents the predicted fold of the ZF domains. Residues at the −1, 2, 3 and 6 positions of each helix contact DNA bases in canonical complexes.^3^ **b)** Homology model built using as templates (Figure S1) the crystal structures of DNA-bound ZBTB38 zinc finger 6 (for ZF1) and Y3 zinc finger of Human YY1 protein (for ZF2); colors as in **a**. **c)** Representative calorimetric titration of ZnCl_2_ into zinc-depleted Loz1AZZ. Top panel, the raw thermogram; bottom panel, integrated heats (black) and best fit to a binding model with two independent sites (red). Best fit parameters are n_1_ = 1.09 ± 0.13, K_d1_ = 7 ± 10 nM, ΔH_1_ = −21 ± 1 kcal mol^-1^, n_2_ = 1.00 ± 0.18, K_d2_ = 290 ± 100 nM, ΔH_2_ = −29 ± 3 kcal mol^-1^.

Given that the Loz1 zinc-responsive domain maps to a region containing two C_2_H_2_-type ZFs and an uncharacterized “accessory domain”, the goal of this study was to investigate whether each of these domains has a role in sensing zinc ions. The sequence of the canonical ZFs of Loz1 are conserved across *Schizosaccharomyces* species, but there is little sequence conservation in the accessory domain.^16^ On the other hand, Ser, His and Asp are known to coordinate zinc ions with modest affinities^23^ which are abundant in the semi-conserved accessory domain (Figure 1a and b). We present calorimetry and heteronuclear NMR studies of the *Spo* Loz1 zinc-responsive region, which show the ZF domains as the likely zinc sensors. NMR data show that the ZFs exhibit dynamics over a range of time scales and bind zinc with differing affinities. In addition, NMR chemical shift analysis and homology modeling provide insights into mechanism of recognition of sequence-specific recognition of LRE DNA. These findings suggest a mechanism by which protein dynamics keep the protein in an inactive state, while zinc binding-coupled folding of Loz1 zinc finger 1 (ZF1) results in specific DNA binding and gene repression.

## Materials and Methods

### Plasmid construction and expression

Loz1AZZ and Loz1ZZ protein constructs were generated by PCR amplifying Loz1 and cloning the resulting PCR products into the *Nde*I and *Bam*HI sites of a pET21a vector (Novagen). Inserts were amplified in 25 μL PCR reactions using *Taq* DNA polymerase (New England Biolabs), Loz1 genomic DNA as the template, and the following oligonucleotide primers: Loz1-REV, 5’-cat<u>ggatcc</u>CTACAAACCATGAATGCGTTGA – 3’, Loz1AZZ, 5’ – cgg<u>catatg</u>TCCAATTATTCTGATCATCAC −3’, and Loz1ZZ, 5’-cat<u>catatg</u>CGCAAAATTGCACAATCCC-3’; restriction sites are underlined, Loz1-template sequences are upper case. Ligated plasmids were transformed into *E. coli* BL21(DE3) competent cells using electroporation and plated onto LB agar plates supplemented with 100 μg/ml of carbenicillin. A single isolated colony was used to inoculate 100 mL of LB broth containing the 100 µg/mL of the antibiotic and grown to an O.D. of 1 at 600 nm. 10 mL of this starter culture was used to inoculate 1 L of M9 minimal medium (6.6 g NaH_2_PO_4_, 3 g of K_2_HPO_4_, 1 g NaCl, 2 mM MgSO_4_, 100 µM CaCl_2_, 1x MEM Vitamin Mix (Gibco), 100 µg/mL carbenicillin) supplemented with 1 g of ^15^NH_4_Cl (Cambridge Isotope Laboratories) and 4 g of D-glucose for U-[^15^N]-protein, or 1 g ^15^NH_4_Cl and 2 g ^13^C-D-Glucose (Martek Isotopes) for U-[^15^N,^13^C]-protein. Cultures for expressing Loz1AZZ were grown at 37 °C to an O.D. 600 of ∼0.8 and expression was induced by addition of 1 mM IPTG supplemented with 100 µM ZnCl_2_. Cells were harvested by centrifugation (4200 x *g* for 10 min at 4°C) four hours after induction. Loz1ZZ expression hampered bacterial growth, therefore cultures were grown at 37°C to an O.D. 600 of 1 to accumulate cell density and were induced overnight at 18°C or 25 °C to reduce cellular metabolic activity. Loz1 point mutants were generated using the pET21a vector encoding Loz1AZZ was as the template for QuikChange site directed mutagenesis (Agilent) using the primers listed in Table S2.

### Protein purification and sample preparation

All buffers were filtered and degassed under vacuum using a bottle top filter prior to use. Dialysis buffers were additionally degassed by bubbling argon through the buffer. Cell pellet from 1 L culture was lysed using sonication on ice in 35 mL Buffer B (50 mM Tris pH 7.5, 1 M NaCl, 10 mM β-mercaptoethanol, 100 μM ZnCl_2_) with half a tablet of cOmplete, Mini EDTA-free Protease Inhibitor Cocktail (Roche, cat# 11836170001) and 0.1% (v/v) Tween-20. The cell debris was pelleted using centrifugation (27,000 x *g* for 45 min at 4 °C) and the soluble fraction containing the protein was diluted 4-fold with Buffer A (50 mM Tris pH 7.5, 10 mM β-mercaptoethanol, 100 μM ZnCl_2_) to reduce salt concentration. The diluted soluble fraction was then loaded at 1 mL/min in Buffer A onto a 5 mL HiTrap SPFF column (GE Healthcare Life Sciences) and eluted at 2 mL/min with a 70 mL gradient to 100% Buffer B; proteins eluted ∼55% Buffer B. The fractions containing the protein were identified from Coomassie-stained 8-16% SDS-PAGE gradient gels (Genscript), pooled and diluted 3-fold with Buffer A. The diluted peak fractions were loaded at 2 mL/min in Buffer A onto a 5 mL HiTrap Heparin column (GE Healthcare Life Sciences) and eluted at 2 mL/min with a 50 mL gradient from 30% to 100% Buffer B; proteins eluted ∼70% Buffer B. The fractions containing the protein were concentrated to 2 mL using centrifugal filters (Millipore-Sigma, 3k MWCO) and then loaded on a Superdex 75 16/60 size exclusion column (GE Healthcare Life Sciences) pre-equilibrated with Buffer S (20 mM Tris pH 7.5, 100 mM NaCl, 2mM TCEP, 100 μM ZnCl_2_, 0.02% NaN_3_). The protein was eluted at 0.75 mL/min and protein fractions were identified using SDS-PAGE. Fractions containing the protein were pooled and concentrations determined using Pierce BCA Protein Assay Kit - Reducing Agent Compatible (Thermo Fischer Scientific, cat# 23250). Integrity, purity and extent of isotopic labeling were assessed by SDS-PAGE with Coomassie staining (Figure S2) and MALDI-TOF mass spectrometry (Bruker Microflex) or electrospray ionization (Q Exactive EMR+, Thermo).

Loz1 point mutants were expressed using the same procedure as with the wild-type protein except for the P480A mutant, which partitioned to the insoluble fraction of the cell lysate. To isolate the P480A Loz1, the insoluble lysis product was washed three times by resuspending the pellet on ice in Wash Buffer (50 mM Tris pH 7.5, 100 mM NaCl, 4 mM TCEP, 2 M urea, and 10% w/v Triton X-100) and pelleted at 27,000g for 30-40 minutes, then one additional time without urea or Triton X-100. The resulting pellet was then resuspended in Extraction Buffer (50 mM Tris pH 7.5, 100 µM ZnCl_2_, 6 M guanidinium HCl, 100 mM NaCl) and centrifuged at 27,000g, 4 °C for 80 minutes. The supernatant was then collected, diluted 20-fold with Buffer A (50 mM Tris, 100 mM salt, 4mM TCEP, 100µM ZnCl_2_) before purifying on 5 mL HiTrap SPFF column as described above.

### Calorimetry

Zinc-free Loz1 was prepared by treatment with a 10-fold molar excess of EDTA over protein, followed by two-step dialysis (500x dilution, each) using 3.5 K MWCO G2 cassettes (Thermo Scientific, cat# 87723) against degassed Buffer H (50 mM HEPES pH 7.5, 100 mM NaCl, 10 mM TCEP) containing Chelex, for at least 12 hours each, at 4°C. Isothermal titration calorimetry was performed using a VP-ITC instrument (Malvern). Zinc solutions were prepared by dilution of zinc chloride stock (10 mM in water) in ITC buffer. Protein concentrations in the ITC cell were 4 to 7 μM, and zinc concentrations in the syringe were 80 to 200 μM. Zinc concentrations were determined using an Agilent Atomic Absorption Spectrometer 240FS. ITC data were recorded in 5 μL first injection followed by 25 injections of 10 μL, spaced 420 seconds apart, at 298K. Origin 7.0 (OriginLab) was used to integrate and fit the data to a binding model with two independent sets of sites to extract the thermodynamic parameters, *n,* Δ*H* and *K_d_* for each site.

### NMR

NMR spectra were recorded at 25°C on 600, 800 or 850 MHz Bruker Avance III spectrometers fitted with 5 mm cryoprobes on samples ranging from 0.5 mM to 1 mM concentration in NMR Buffer (20 mM Tris pH 7.5, 100 mM NaCl, 2 mM TCEP, 100 μM ZnCl_2_, 0.02% NaN_3_, 10% (v/v) ^2^H_2_O and 200 μM DSS as an internal standard; heteronuclear shifts were referenced indirectly to DSS from the corresponding gyromagnetic ratios). Data were processed using NMRpipe^24^ or NMRFx^25^ and analyzed using NMRViewJ^25^.

### Resonance Assignments

Data from 3D HNCO, HNCA, HNCOCA, HNCACB, CBCACONH experiments were used to assign backbone H^N^, N, C’, C^α^ and C^β^ atoms. 3D HCCH-TOCSY, HCCH-COSY, H(CCO)NH-TOCSY and (H)CC(CO)NH-TOCSY spectra were used to obtain additional sidechain proton and carbon assignments, respectively. The chemical shift index for C_α_ and C_β_ were obtained from *TALOS-N* software^26^. Chemical shift perturbations (CSPs) were calculated using the amide chemical shifts according to:

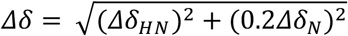

Where Δδ is the compounded chemical shift perturbation, Δδ _HN_ is the chemical shift perturbation of the amide proton, Δδ _N_ is the chemical shift perturbation of the amide nitrogen.

### NMR relaxation

^1^{H}-^15^N heteronuclear NOE, *R*_1_ and *R*_1_ρ data were recorded at 600 and 850 MHz for Loz1AZZ, and at 800 MHz for Loz1ZZ, following published methods^27^. *R*_1_ and *R*_1_ρ data were acquired in an interleaved fashion with relaxation delays of 40, 200 (x2), 520, 720 (x2) ms and 2, 34, 68 (x2), 100 (x2) ms, respectively using a recycle delay of 2 s. ^1^{H}-^15^N hetNOE values were obtained from ratios of signal intensities for spectra collected in presence and absence of ^1^H pre-saturation during a 4 s recycle period. The relaxation rates were obtained by fitting peak intensities to an exponential decay function.

^15^N relaxation dispersion CPMG data were recorded at 600 and 800 MHz proton frequencies for each Loz1AZZ and Loz1ZZ construct following published methods^28^. The experiments on both instruments were acquired using a recycle delay of 2 s and varying CPMG pulse train frequency: for 800 MHz data, 82.6 μs pulses in pulse train of 50 ms with 19 points (0, 20, 40 (x2), 60, 80, 100, 120, 160, 200, 280, 440, 600, 760, 920, 1080, 1240 (x2), 1400 Hz); and for 600 MHz data, 87.2 μs pulses in a pulse train of 40 ms with 21 points (0, 25, 50(x2), 75, 100, 125, 150, 175, 200, 250, 300, 350, 400,500, 600, 700, 800, 900(x2), 1000 Hz). Dispersion data from both 600 and 800 MHz were fit simultaneously with the Carver-Richards two-state exchange model for individual residues and globally for groups of residues, using GUARDD^29^. Good quality fits had small and random residuals.

### NMR-monitored zinc titrations

Zinc-free protein was generated by treatment with 10-fold molar excess EDTA over protein, followed by overnight dialysis (500x dilution) using 3.5K MWCO G2 cassettes (Thermo Scientific, cat# 87723) against degassed NMR-buffer without ZnCl_2_, at 4 °C. Zinc-coupled binding/folding of apo Loz1AZZ construct was monitored by recording 2D {^1^H}-^15^N HSQC spectra over a range of added zinc concentrations. 10 mM zinc chloride solution in NMR buffer, made by diluting 1 M zinc chloride stock (in water), was titrated into 200 μM [U-^15^N]-Loz1AZZ and incubated at room temperature for 15 minutes before data collection. Fractional peak intensities were calculated from the ratio of each peak intensity to the maximum peak intensity measured from all {^1^H}-^15^N-HSQC experiments for that residue.

### DNA substrate and DNA-protein complex preparation

ssDNA oligos: 5’-d(GCGACGATCA TTGG)-3’ and 5’-d(CCAA TGATCGTCGC)-3’ (IDT, Inc.) were resuspended in DNA buffer (20 mM Tris pH 7.5, 50 mM NaCl) and annealed to obtain dsDNA substrate by heating equimolar mixture of oligos to 95 °C in a water bath for 15 minutes and cooling in water bath overnight to room temperature. One dimensional ^1^H spectra revealed eight imino signals (4 G-C and 4 A-T pair). PD-10 column (GE Healthcare Life Sciences) was used to exchange the dsDNA into NMR buffer. Concentration of DNA was measured using UV-Vis spectrometry absorbance at 260 nm. 2 mM of purified Loz1AZZ in NMR buffer was titrated into 130 μM DNA substrate to reach an equimolar concentration.

### Homology modeling and docking

Each Loz1 zinc finger (ZF) domain was individually submitted for homology modeling using the SwissModel webserver.^30^ The template (Figure S1) for modeling ZF1 was ZF 1 of ZBTB38 (residues 1010 - 1033) bound to DNA (PDB:6E93), and ZF2 was modeled using ZF 3 of YY1 (residues 349 – 377) bound to DNA (PDB: 1UBD).^31,32^ The resulting models were superposed onto zinc finger 2 and 3 of the YY1-DNA complex using the align function of PyMOL (Schrödinger). The LRE DNA model was generated by first using the X3DNA webserver to obtain the helical parameters from the YY1 DNA substrate, and then rebuilding the LRE consensus sequence by mutating positions C6, C9, and T12 of the YY1 substrate to G6, G9, and C12 of the LRE using the X3DNA webserver in conjunction with previously obtained DNA helical parameters.^33^ After superposing the two zinc fingers of Loz1 onto the modeled LRE, the chain break present from homology modeling each finger independently was closed using the kinematic closure protocol in Rosetta.^34^ Knowledge of canonical zinc finger recognition of DNA substrates was used to generate a set of ambiguous and unambiguous restraints from Loz1 ZZ to the LRE DNA substrate, including restraints to maintain correct geometries for coordinating zinc in each ZF.^35^ These restraints and homology models were submitted to the HADDOCK 2.4 webserver for docking and the lowest energy model from the best scoring cluster (RMSD, HADDOCK Energy, and restraint energy violation) was selected as the Loz1-LRE complex model for analysis.^36^

## Results

### Loz1 zinc finger domains exhibit differential affinity towards zinc

To characterize the stoichiometry and thermodynamics of zinc binding to Loz1 we used isothermal titration calorimetry (ITC) to measure heat evolved upon titrating zinc into zinc-depleted protein. Loz1 constructs containing (Loz1AZZ) or lacking (Loz1ZZ) the accessory domain were recombinantly expressed in *E. coli* BL21(DE3) cells and purified in buffers supplemented with zinc (100 μM). Zinc-free “apo-Loz1” was obtained by treatment with EDTA followed by dialysis against Chelex-treated, degassed buffers containing TCEP as a reducing agent to prevent cysteine oxidation. ITC titrations of zinc chloride solutions into apo Loz1AZZ resulted in biphasic thermograms that could be fit by a model with two independent sites (Figure 1c). The zinc affinity for the first binding event was found to be in the nanomolar range (*K*_d1_ = 7 ± 10 nM), whereas the affinity for the second zinc binding event was 1.6 orders of magnitude weaker (*K*_d2_ = 300 ± 100 nM). ITC experiments with the construct lacking the accessory domain (Loz1ZZ, res. 461-522) yielded similar thermograms and affinities as those for Loz1AZZ (K_d1_ = 4 ± 6 nM and K_d2_ = 350 ± 90 nM; Figure S3). These results suggest that zinc sensing by Loz1 is accomplished by differential zinc binding by the two zinc fingers, with no apparent influence of the accessory domain on the binding affinities. There was also no evidence of zinc binding to the accessory domain.

### NMR spectra reveal a disordered accessory domain and slow *cis-trans* X-Pro peptide bond isomerization in the ZF1-ZF2 linker

Heteronuclear NMR spectroscopy provided evidence for conformational heterogeneity in the Loz1 zinc-responsive element. Loz1AZZ was uniformly labeled with ^15^N, or ^13^C and ^15^N and purified to homogeneity as evidenced by electrospray ionization mass spectrometry under native conditions, which revealed a single species with a mass corresponding to Loz1AZZ bearing > 99% ^15^N enrichment and two bound zinc atoms (Figure S2). Despite chemical and compositional homogeneity, 2D ^1^H-^15^N correlation spectra of zinc-bound Loz1AZZ (426-522) featured doubling of several of the resonances, with minor peak intensities of approximately 20% relative to the major peaks (Figure 2a). In addition, far fewer amide resonances were observed than would be expected for expected for a 97-residue protein, and many resonances were much broader than would be expected for the small 11.2 kDa protein construct. At lower temperatures (5 °C), additional broad signals could be observed in the random coil region of the spectrum (Figure S4), while resonance doubling persisted. Lastly, amide resonances exhibited highly variable intensities and line widths, indicative of exchange broadening.

**Figure 2.**
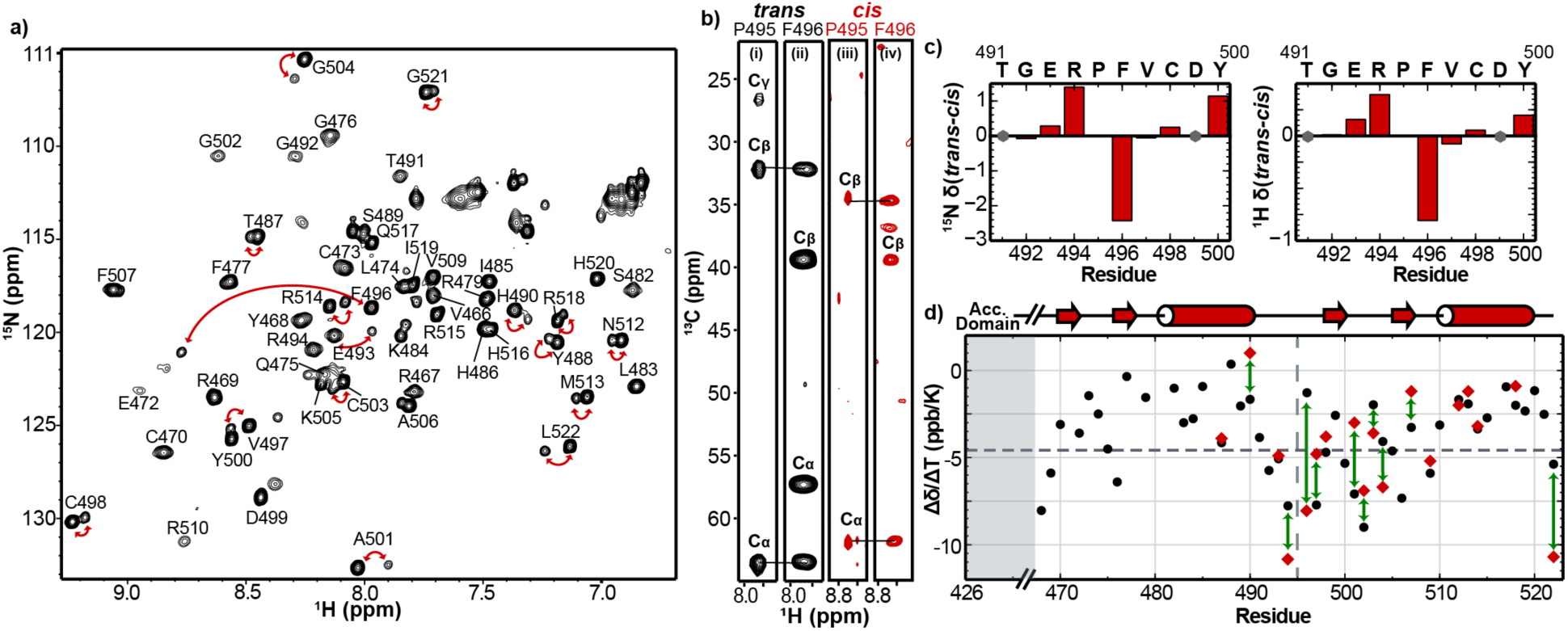
The Loz1 zinc responsive element exhibits an unstructured accessory domain and slow exchange due to *cis-trans* isomerization in the *TGERP* linker. a) Two-dimensional ^1^H-^15^N HSQC spectrum of Loz1AZZ with assigned backbone amide resonances from the zinc finger region. Doubling of signals is observed for residues in ZF2 and helical region of ZF1 (red arrows). Absence of signals from the accessory domain indicates it is poorly structured. b) Strips from 3D CC(CO)NH-TOCSY (i and iii) and HNCACB (ii and iv) spectra highlighting C^α^ and C^β^ chemical shifts of Pro495 and Phe496 residues in the *trans* and *cis* conformers. c) Chemical shift differences (δ*_trans_* − δ*_cis_*) between the *trans* and *cis* Arg494-Pro495 conformations for affected amide proton and nitrogen resonances in Loz1AZZ. Residues for which only one signal was observed are marked with a grey circle. These data indicate that structural effects of *cis-trans* isomerization extend beyond the ZF1-ZF2 linker. d) Amide proton temperature coefficients reveal differences in solvent exposure of backbone amides *trans* (black circles) and *cis* (red diamonds) conformations of the Arg494-Pro495 peptide bond.

Conventional double- and triple resonance NMR experiments enabled us to assign ∼95% of the resonances observed in two-dimensional ^1^H-^15^N HSQC spectra (Figure 2a). Most of the resonances could be assigned to the two ZFs, while no resonances were assigned to residues in the accessory domain. Backbone resonance assignments included 50 out of 55 (91%) amides in the zinc finger and linker region, 80% of the C’ and 95% of the C^α^ resonances. Some residues from predicted loop regions of the zinc fingers (T471, S478, S511 and N508) and S481 from ZF1 helix could not be assigned, likely due to exchange broadening. Three dimensional heteronuclear TOCSY-type experiments (Table S1) were used to obtain additional sidechain proton and carbon assignments and 80% of aliphatic protons from the zinc finger residues could be assigned. Secondary chemical shifts were generally consistent with the predicted ββα fold for each zinc finger (Figure S5). NMR spectra of the construct lacking the accessory domain (Loz1ZZ) were nearly superimposable on those of Loz1AZZ (Figure S3), with differences limited to small perturbation of resonances from adjacent residues Val466 and Arg467, and the absence of two unassigned signals. These observations indicate that the accessory domain exhibits intermediate time scale disorder and is dispensable for structural integrity of the zinc fingers and for their zinc-sensing function.

Doubling of the amide resonances results from slow *cis*-*trans* isomerization of Arg-Pro peptide bond in the TGERP linker between the two zinc finger domains. Doubled resonances mapped to the end of the ZF1 helix, the TGERP^495^ linker, and many residues in ZF2. The chemical shift difference between C^β^ and C^γ^ of Pro^495^ is consistent with a *trans*-conformation of the major state; the minor state Pro C^γ^ resonance was not observed, but the C^β^ chemical shift is comparable to that for *cis*-conformation^37,38^ (Figure 2b). This assignment is substantiated by the significant upfield shift in ^1^H and ^15^N resonances of Arg494 (i−1 residue) in the *cis*-state (Figure 2c). Further, the upfield shift (δ_trans_ - δ_cis_ ∼ 1.6 ppm) of C^α^ resonance for the i-2 residue is similar in sign and magnitude to the C^α^ chemical shift change (δ_trans_ - δ_cis_ ∼1.2 ppm) of the isomerizing Pro, diagnostic of the *cis* state.^39^ To support the conclusion that X-Pro isomerization at Pro495 is responsible for the observed signal doubling we prepared samples in which each of the two proline residues in Loz1AZZ was replaced with Ala: P495A and P480A. Two-dimensional NMR spectra of P480A-Loz1 exhibited the same signal doubling, while only one set of signals is observed for the P495A variant (Figure S6). Although canonical DNA-binding C_2_H_2_ ZF proteins feature TGE(R/K)P linkers^4,40^, slow *cis-trans* isomerization has not previously been reported for this class of proteins. Substitution of Arg494 in the linker with Lys also did not eliminate the slow exchange behavior (Figure S7). No exchange cross-peaks were observed in ZZ exchange^41^ spectra probed up to 900 ms, consistent with exchange rates much slower than 1 s^-1^.^42,43^

NMR spectra of major and minor states reveal that structural perturbations from *cis-trans* isomerization of the Arg494-Pro495 peptide bond are not limited to the linker. Amide chemical shift perturbations from *cis-trans* isomerization (i.e., signal doubling) are observed throughout ZF2 to the C-terminal residue Leu522 (Figure 2c, S8). We measured amide proton temperature coefficients (Δδ/ΔT in ppb K^-1^) by recording ^1^H-^15^N HSQC spectra at five-degree intervals from 278 to 308 K (Figure S9); these report on secondary structure stability via the hydrogen bond strength and its thermal expansion, with smaller Δδ/ΔT being associated with stronger intramolecular hydrogen bonds, whereas the largest negative Δδ/ΔT are associated with hydrogen bonding to solvent^44–46^. Temperature coefficients of amide protons in the *cis* and *trans* species (Figure 2d) show the largest differences for residues in the β-hairpin region of ZF2, consistent with their proximity; however, just as many amides experience a positive and negative change in Δδ/ΔT. These data indicate that *cis-trans* isomerization of the X-Pro peptide bond in the TGERP linker has structural consequences on the entirety of ZF2, not just the linker.

### ZF1 has lower zinc affinity than does ZF2

To identify the zinc finger domain with weaker zinc affinity, making it a candidate for direct zinc sensing, we performed NMR-based zinc titrations (Figure 3). [U-^15^N]-Loz1AZZ was stripped of bound zinc using EDTA as for ITC experiments. The ^1^H-^15^N HSQC spectrum of zinc-free Loz1AZZ exhibited a high degree of line broadening, with few well-resolved signals (Figure 3b, left). Most signals cluster in a narrow range of ^1^H frequencies of 8 – 8.5 ppm, with little signal dispersion in both dimensions, characteristic of aggregation in the sample or conformational heterogeneity on a ms-μs time scale^47^. Titrating zinc into the protein solution resulted in the incremental appearance of signals corresponding to the zinc-bound, folded form of the protein, while broad resonances from the apo form diminished (Figure 3b, center and right). At one equivalent of added zinc, signals from ZF2 residues were fully populated (Figure 3c). Signals from ZF1 did not completely saturate even after addition of two molar equivalents of zinc. These data indicate that ZF1 has the lower affinity zinc site identified by ITC (Figure 1).

**Figure 3.**
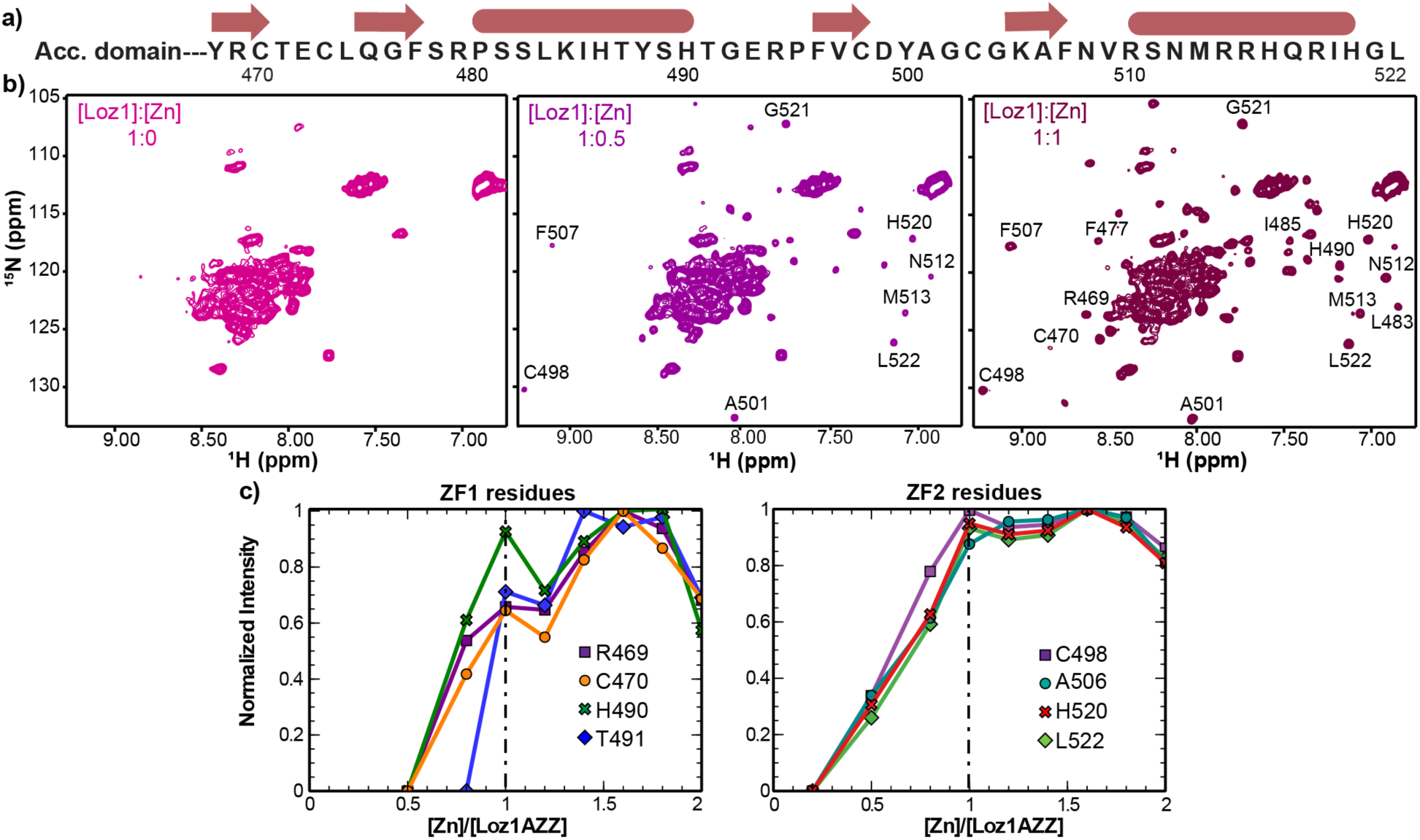
Loz1 ZF1 coordinates zinc with weaker affinity. **a)** Loz1AZZ amino acid sequence with a schematic representing the predicted secondary structure. **b)** ^1^H-^15^N HSQCs of Loz1AZZ in zinc-free apo form (left), 0.5 equivalents of zinc (middle), and equimolar zinc (right). Apo-Loz1AZZ amide backbone amide signals are clustered around 8.3 ppm, diagnostic of a disordered protein. Upon addition of limiting zinc, resonances from folded ZF2 appear in the spectrum whereas ZF1 resonances appear upon titration with additional zinc. **c)** Representative plots of normalized intensities of residues from ZF1 and ZF2 against [Zn]:[Loz1AZZ] molar ratio. The dashed line corresponds to equimolar zinc to protein ratio (third panel in b)) at which point ZF2 intensities saturate while ZF1 intensities continue to increase, indicating weaker zinc binding to ZF1.

### Loz1 exhibits heterogeneity on fast and intermediate time scales

Doubling of NMR signals due to *cis-trans* isomerization revealed structural heterogeneity on slow time scales ( > 1 s), while non-uniform intensities and broad lines were indicative of internal motions on intermediate time scales (µs-ms). We first probed fast (ps-ns) time scale dynamics of the major state of Loz1AZZ by recording ^15^N spin relaxation rates *R*_1_, *R*_1ρ_ and {^1^H}-^15^N heteronuclear NOE spectra; *R*_2_ values were computed from *R*_1_-corrected *R*_1ρ_ values (Figure 4; Figure S10). The low signal-to-noise for the data from the *cis* state precluded its analysis. Heteronuclear NOE values for *trans* Loz1AZZ were generally uniform for residues in the predicted helices, with values near the 10% trimmed-mean for the entire dataset of 0.63 at 600 MHz, excluding linker residues. Values decreased as far as 0.44 for residues in the linker and the zinc binding loops, indicating increased ps-ns flexibility for those regions. The *R*_1_/*R*_2_ ratio was found to differ between ZF1 and ZF2, suggestive of anisotropic tumbling^48–51^. However, similar data collected on Loz1ZZ exhibited similarly elevated *R*_2_ values for ZF1, despite the absence of the accessory domain, suggesting that the elevated *R*_2_ arises from exchange broadening by an underlying dynamic process at the µs-ms timescale.

**Figure 4.**
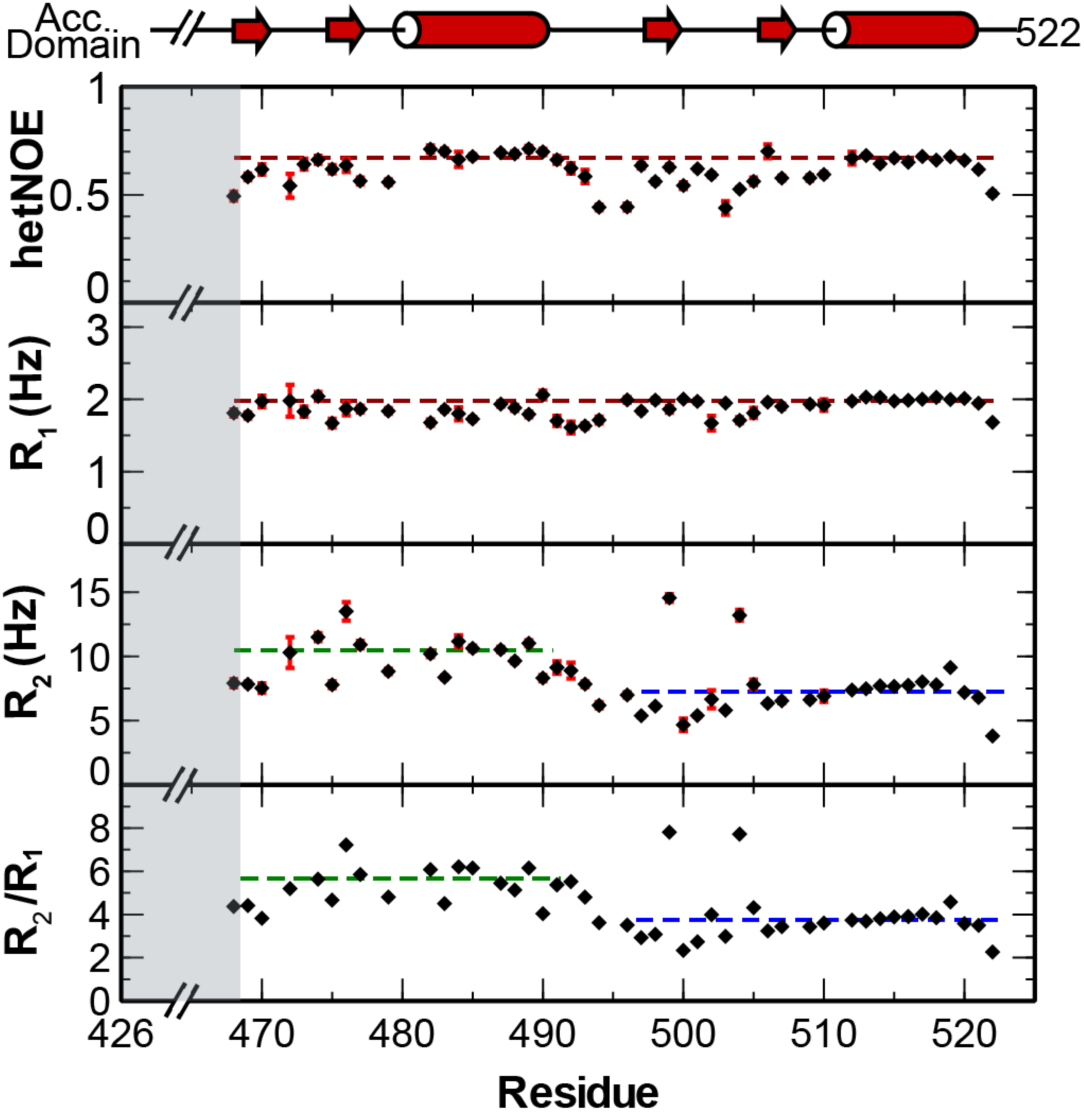
^15^N relaxation data reveal elevated *R*_2_ values for ZF1. Loz1AZZ {^1^H}-^15^N heteronuclear NOE, *R*_1_, *R*_2_ and *R_2_/R_1_* ratios from data recorded at 600 MHz. Error bars for the heteronuclear NOE data were obtained from peak height uncertainties based on average noise levels in the NMR spectra. Uncertainties in *R_1_* and *R_2_* values were obtained from replicates. Dashed lines represent the 10% trimmed-mean values excluding linker region. The hetNOE and *R_1_* values could be described by a uniform trimmed mean value, whereas ZF1 and ZF2 exhibited different mean *R*_2_ values.

Amide ^15^N Carr-Purcell Meiboom-Gill relaxation dispersion (CPMG RD)^28^ experiments at two magnetic fields revealed intermediate exchange dynamics in Loz1AZZ. Indeed, the effective transverse relaxation rate *R*_2,eff_ of many amide resonances were found to depend strongly on the rate at which the CPMG refocusing pulses were applied (i.e., ν_CPMG_) (Figure 5). These dispersion curves, with exchange contributions to transverse relaxation *R*_ex_ as high as 13 s^-1^, mapped to all assigned amides in ZF1 and the linker region, as well as Asp499 and Gly504 in ZF2 (Figure 5a). Weak (< 2 s^-1^), or no dispersions were observed in most of ZF2. Similar relaxation dispersion profiles were observed for the same residues for Loz1ZZ, indicating that these dynamics are an inherent to ZF1, not a consequence of transient interactions with the accessory domain (Figure S10).

**Figure 5.**
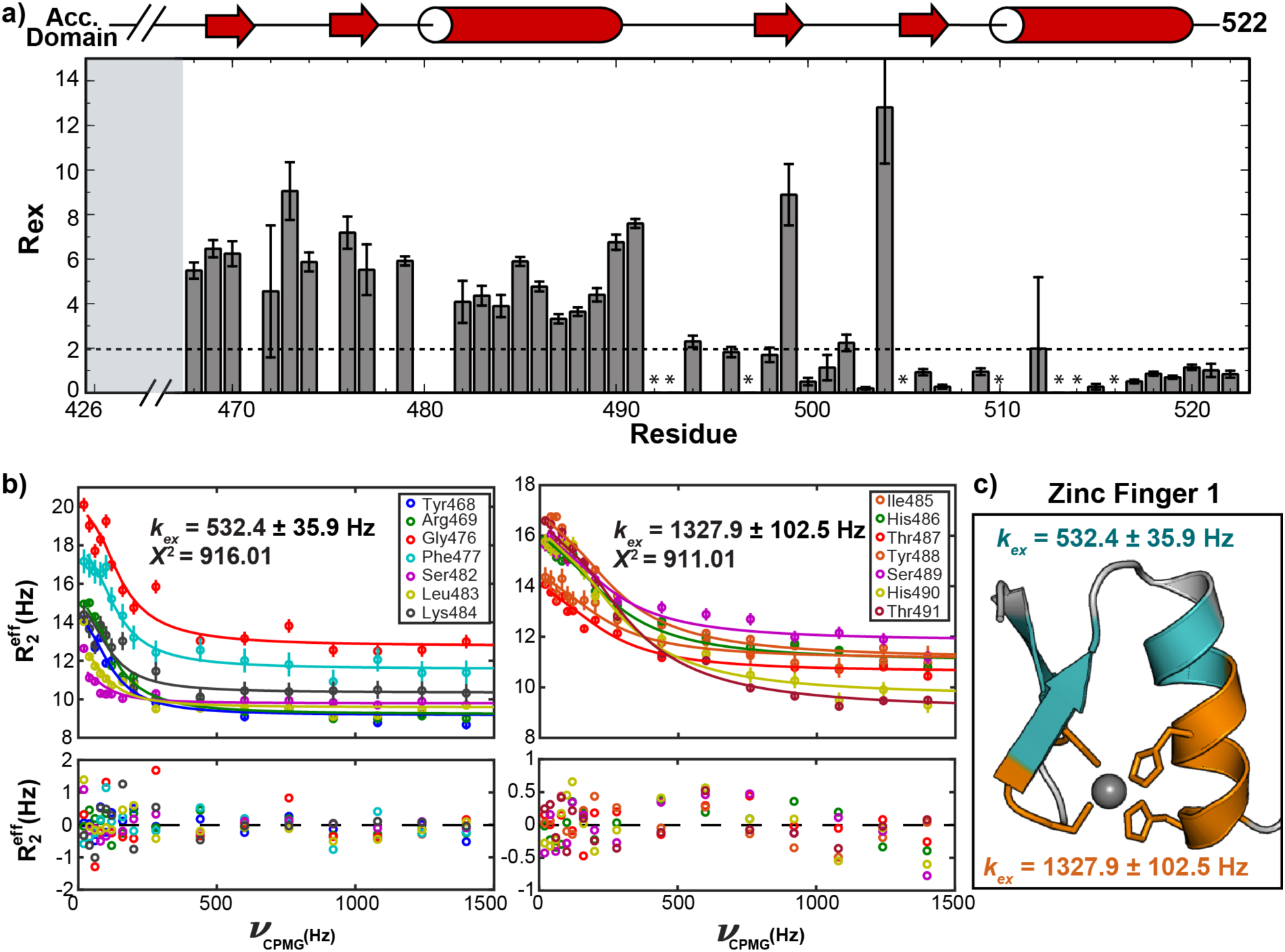
Loz1 ZF1 undergoes complex dynamics at µs-ms time scales. a) Exchange contributions to transverse relaxation *R*_ex_ for Loz1AZZ, recorded at 800 MHz, obtained by two-field fitting of CPMG dispersion curves for each residue with the Carver-Richards two-site exchange model. An asterisk identifies assigned residues for which good fits of the dispersion curves could not be obtained, or had *R*_ex_ values close to zero; the dashed line is the cut-off below which dispersions were considered unreliable. **b**) Left, global fit of a group of residues undergoing “slow” exchange with a *k*_ex_ of 532 s^-1^. Right, global fit of residues undergoing “fast” intermediate exchange, with a fitted *k*_ex_ of 1,328 s^-1^. **c)** Homology model of ZF1 with “slow” exchanging residues in blue, and “fast” exchanging residues in orange. Residues that are either unassigned or did not yield reliable fits to a two-state exchange model are grey.

Relaxation dispersions in Loz1AZZ could not be explained by concerted exchange between two well-defined states. Global fitting of dispersion curves for ZF1 with the Carver-Richards two-state exchange model^29^ resulted in poor fits and large residuals. However, residues could be separated into two groups, each of which could be fit to a global two-site model – the region close to the zinc coordinating region of the ZF with faster exchange rate (*k*_ex_ = 530 ± 40 s^-1^) and slower exchange rate (kex = 1330 ± 40 s^-1^) for residues away from the zinc coordinating site. (Figure 5b and c). This behavior is consistent with two independent motional modes, or perhaps exchange between more than two states, the fitting of which is beyond the precision of the available data and Carver-Richards approach^52^. These observations highlight a strong parallel between weak zinc affinity and intermediate time scale conformational exchange.

### DNA-bound Loz1AZZ exhibits a single conformation

To test whether the *cis-trans* isomerization observed in free Loz1AZZ persists when bound to DNA, 2D ^1^H-^15^N NMR spectra were recorded in the presence of a 14 base pair DNA duplex containing the LRE element (5’-GC**<u>G</u>**<u>ACG**ATC**</u>ATTGG-3’). The Loz1AZZ-LRE complex was assembled in NMR buffer by titrating [U-^15^N]-Loz1AZZ into the dsDNA. Upon formation of the DNA complex, chemical shift perturbations were observed throughout the spectrum (Figure 6a), consistent with an extensive protein-DNA surface and remodeling of the inter-finger interface^53^. Increased uniformity of amide signal intensity is consistent with quenching of µs-ms exchange upon binding to DNA. Doubling of amide resonances was also not observed, consistent with a canonical *trans* Arg-Pro peptide bond conformation in the protein-DNA complex. As in the absence of DNA, fewer than the 95 protein amide resonances were observed, indicating that the accessory domain does not adopt a well-defined structure when bound to an LRE. Homology modeling of the two zinc finger domains and docking to a model DNA duplex (Figure 6b) places residues at the recognition positions −1, 2, 3 and 6 in position to recognize the consensus nucleotides in the LRE with Ile485 and Val509 packing in the inter-finger and DNA interface.

**Figure 6.**
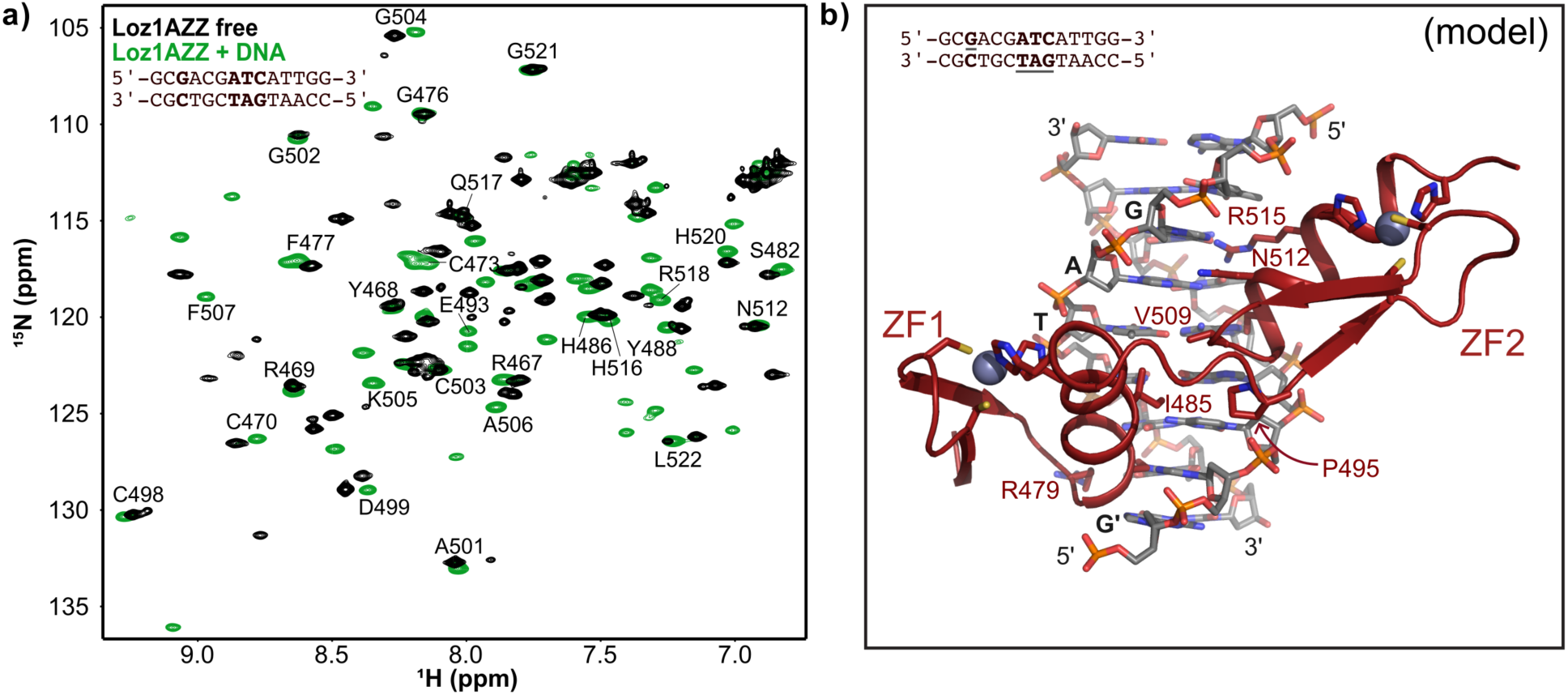
DNA binding quenches both fast and slow dynamics in Loz1AZZ. **a)** Overlay of ^1^H-^15^N HSQC spectra Loz1AZZ in absence (black) and presence (green) of near equimolar LRE DNA. The DNA substrate sequence is shown with the consensus nucleotides in bold. Indicated amide assignments in the bound state are inferred from proximity to isolated signals in spectra of the free protein. Absence of signal doubling indicates the *cis* Arg-Pro conformer is no longer populated, while more uniform peak shapes are indicative of reduced motion on the µs-ms time scale. b) Model of zinc-loaded Loz1AZZ bound to the consensus DNA. Side chains of residues at the −1, 2, 3 and 6 positions of each helix are shown as sticks, and the consensus base positions in the bottom strand are indicated in bold.

## Discussion

We used NMR and binding assays to characterize the domains of Loz1 that are sufficient for its zinc-responsive function. Calorimetric and NMR titrations localized the zinc-sensing function to the first C_2_H_2_ zinc finger in Loz1. ZF1 binds zinc weakly (*K*_D_ ∼300 nM) (Figure 1, Figure 3) and exhibits complex dynamics on the timescale of 10^-6^ – 10^-3^ that leads to line broadening and relaxation dispersion in NMR experiments (Figure 5)^52^. Motions on these timescales are relevant to processes that may enable non-concerted binding and release of zinc, such as due to histidine switching^54^ or ligand flipping^55^. These atypical properties reveal a mechanism for zinc-responsive gene repression wherein elevated cellular zinc levels promote zinc-dependent folding of ZF1. The structured sensory finger can then make base-specific and backbone interactions with LRE DNA sequence, in addition to those contributed by the structural finger (ZF2).

In addition to the dynamic properties that make ZF1 a sensor, we found evidence for slow *cis-trans* isomerization of the Arg-Pro peptide bond in the TGERP linker between the two zinc fingers. Low-level population of a *cis* conformer is not unusual in flexible polypeptides containing Pro residues ^39,42^ but this behavior has not previously been noted for other C_2_H_2_ zinc finger proteins with canonical TGE(K/R)P linkers.^56^ NMR studies on classical C_2_H_2_ zinc finger proteins have generally found the ZF domains to not interact significantly in the absence of DNA, but because of restraints imposed by the short linkers tumble anisotropically, with correlated diffusion^51,57,58^. In the major *trans* conformation of Loz1AZZ the zinc-fingers exhibit different *R*_2_/*R*_1_ ratios, consistent with lack of inter-finger interactions. We estimate the population of the *cis*-proline conformation to be ∼15-20% based on peak intensities, which is higher than what it typically found in model peptides and unfolded proteins.^39,42^ When bound to DNA, only a single conformation of Loz1 is observed, consistent with the *trans* conformation observed with other such protein-DNA complexes.^3,4,31,32,59,60^

While the biological significance of the *cis-trans* X-Pro isomerization in the Loz1 TGERP linker has yet to be examined, evidence from other systems suggest a potential role in regulating zinc-activated repression^17,61,62^. Post-translational modifications in canonical TGE(R/K)P linker sequences, such as Thr phosphorylation during mitosis in Ikaros, Sp1, ^63^ and Yin Yang 1 (YY1) protein, or acetylation of Lys in YY1^64,65^ or erythroid Krüppel-like factor (EKLF), have been associated with modulation of DNA-binding activity.^66^ In cells, peptidyl propyl cis/trans isomerases catalyze the interconversion of the cis- and trans-forms of peptide bonds, often in response to specific environmental or cellular signals^67,68^. As these regulated conformational changes can affect protein folding, stability and activity, *cis-trans* isomerization of X-Pro bonds can function as a molecular switch to control DNA binding, subcellular localization, enzymatic activity, homo-dimerization, and protein-protein interactions^61,68,69^. As alterations in the ratio of *trans* to *cis* forms would potentially influence the levels of Loz1 available to bind to DNA under a given growth condition, X-Pro *cis-trans* isomerization in Loz1 presents a potential additional layer for regulated zinc finger function. Future studies will be required to test this hypothesis and to determine whether this might be a general mechanism to modulate the activity of zinc finger proteins with TGE(R/K)P linkers.

Analysis of the Loz1-LRE docking model (Figure 6) contributes to an expanded understanding of the diversity of DNA sequence recognition mechanisms employed by C_2_H_2_ ZF proteins.^70^ The model is overall consistent with a dominant role for residues at the −1, +3 and +6 positions of each α-helix (Figure 1) in direct readout of the DNA sequence. Prior ChIP-Seq and in vivo reporter assay experiments identified a consensus of Loz1 DNA recognition element (LRE) of 5’-GnnGATC-3’.^20^ In the docking model, the sidechain guanidinium of Arg479 at the −1 position of ZF1 is positioned near the 5’ G of this sequence where it could participate in Hoogsteen-face recognition of the N7 and O6. In ZF2, Asn512 at the +3 and Arg515 at the +6 position are positioned for canonical recognition of the GA step on the complementary strand, 5’-GATCnnG-3’, while a hydrophobic contact between Ile485 (ZF1 +6 position) and Val509 (ZF2 −1 position) coincide with the C5 methyl group of T on that complementary strand. ZF1 has Ser482 at position +3 where it could make favorable but perhaps not highly selective interactions major groove. While these structural features await experimental validation, the correspondence between a naïve homology model and the known sequence preference suggests that major features of the model may be correct and provide insights into the mechanism for site selection by Loz1.

Deletion experiments established that the Loz1 accessory domain (residues 426-460) is part of the minimal construct required for Loz1-mediated transcriptional repression of *zrt1* and *adh4* in vivo.^21,22^ The results reported here indicate that it does not appear to function significantly in zinc sensing or site recognition in vitro. Similarity in NMR spectra and zinc binding affinities for AZZ and ZZ constructs indicates that the accessory domain does not become structured upon binding zinc, nor does it alter the affinity of stoichiometry of zinc binding. In the yeast regulatory protein ADR1, an additional 20-residue proximal accessory region becomes structured upon DNA binding and contributes to complex stability,^71^ whereas crosslinking experiments with the *Drosophila* protein Tramtrack reveal that a similar domain feature contributes to both specific and non-specific DNA binding.^72^ In mammalian MTF-1, ZF5 and ZF6 are dispensable for zinc-induced DNA binding activity and reporter gene activation but are necessary to trigger chromatin-remodeling and activation of the native MT-1 promoter.^73^ Absent a role in zinc sensing, the Loz1 accessory domain function may instead play a role in stabilizing the DNA bound state or facilitating target gene repression within cells.

The atypical properties of the Loz1 zinc finger domains may provide insight into how the activity of other regulatory factors can be altered in response to cellular zinc status. The human protein ZNF658 was shown to inhibit transcription of multiple genes and act on rRNA involved in ribosomal biogenesis, in response to cellular zinc levels.^74^ This protein contains 21 zinc finger domains. Zinc finger linker sequences have been shown to play a role in zinc-sensing in MTF-1. Those studies showed that replacement of a non-canonical RGEYT linker between zinc fingers 1 and 2 with a canonical TGEKP linker resulted in loss of zinc sensing and constitutive binding of mouse MTF-1 to its target DNA.^75^ Another zinc finger containing protein that may be regulated by zinc occupancy is the APC/C inhibitor EMI2, which is part of the cytostatic factor complex that helps maintain the mature, unfertilized mammalian egg in metaphase II arrest ^76^. During fertilization, transient increases in calcium trigger the rapid expulsion of zinc from the cell, which in turn leads to egg activation and the resumption of meiosis. As this event is dependent upon EMI2, changes in the occupancy of its zinc binding domains could directly coordinate meiotic progression of the mammalian egg.

In summary, our observations reveal a mechanism for Loz1 zinc-responsive gene repression and provide insight into properties of zinc finger domains that enable them to have a sensing function. In eukaryotes, the importance of zinc as a regulator of cellular function has been well established.^77,78^ In addition to homeostasis mechanisms that maintain optimal zinc levels, in certain cells types dynamic zinc signals, zinc waves, and zinc fluxes can result in rapid and transient changes cellular zinc concentrations. Since all these processes have the potential to affect the activation of various zinc binding proteins, knowledge of properties that lead to differences in affinities of zinc binding sites may help identify other proteins and cellular processes whose activity is regulated by zinc.

## ACKNOWLEDGEMENTS

OSU CCIC staff members Alex Hansen and Chunhua Yuan assisted NMR data acquisition; Antonia Duran contributed to sample preparation. Melody (Pepsi) Holmquist, William Moeller, Vicki Wysocki (OSU) and her lab members provided support for ESI MS experiments. Stevin Wilson and Carlos Cardona-Soto and other members of the Bird and Foster lab provided stimulating discussion.

## ACCESSION CODES

*Schizosaccharomyces pombe* Loz1, GenBank: CAB61785.2, PomBase ^79^: SPAC25B8.19c

## References

1. Chasapis CT, Spiliopoulou CA, Loutsidou AC, Stefanidou ME (2012) Zinc and human health: An update. Arch. Toxicol. 86:521–534.

2. Cassandri M, Smirnov A, Novelli F, Pitolli C, Agostini M, Malewicz M, Melino G, Raschellà G (2017) Zinc-finger proteins in health and disease. Cell Death Discov. 3:17071.

3. Wolfe SA, Nekludova L, Pabo CO (2000) DNA recognition by Cys2His2 zinc finger proteins. Annu Rev Biophys Biomol Struct 29:183–212.

4. Nagaoka M, Nomura W, Shiraishi Y, Sugiura Y (2001) Significant Effect of Linker Sequence on DNA Recognition by Multi-Zinc Finger Protein. Biochem. Biophys. Res. Commun. 282:1001–1007.

5. Krizek BA, Merkle DL, Berg JM (1993) Ligand variation and metal ion binding specificity in zinc finger peptides. Inorg. Chem. 32:937–940.

6. Laity JH, Lee BM, Wright PE (2001) Zinc finger proteins: new insights into structural and functional diversity. Curr. Opin. Struct. Biol. 11:39–46.

7. Potter BM, Feng LS, Parasuram P, Matskevichi VA, Wilson JA, Andrews GK, Laity JH (2005) The six zinc fingers of metal-responsive element binding transcription factor-1 form stable and quasi-ordered structures with relatively small differences in zinc affinities. J. Biol. Chem. 280:28529–28540.

8. Bird AJ (2003) Zinc fingers can act as Zn2+ sensors to regulate transcriptional activation domain function. EMBO J. 22:5137–5146.

9. Choi S, Bird AJ (2014) Zinc’ing sensibly: controlling zinc homeostasis at the transcriptional level. Metallomics 6:1198–1215.

10. Heuchel R, Radtke F, Georgiev O, Stark G, Aguet M, Schaffner W (1994) The transcription factor MTF-1 is essential for basal and heavy metal-induced metallothionein gene expression. EMBO J. 13:2870–2875.

11. Chen X, Agarwal A, Giedroc DP (1998) Structural and functional heterogeneity among the zinc fingers of human MRE-binding transcription factor-1. Biochemistry 37:11152–11161.

12. Potter BM, Feng LS, Parasuram P, Matskevichi VA, Wilson JA, Andrews GK, Laity JH (2005) The six zinc fingers of metal-responsive element binding transcription factor-1 form stable and quasi-ordered structures with relatively small differences in zinc affinities. J. Biol. Chem. 280:28529–28540.

13. Chen X, Agarwal A, Giedroc DP (1998) Structural and functional heterogeneity among the zinc fingers of human MRE-binding transcription factor-1. Biochemistry 37:11152–11161.

14. Guerrerio AL, Berg JM (2004) Metal Ion Affinities of the Zinc Finger Domains of the Metal Responsive Element-Binding Transcription Factor-1 (MTF1). Biochemistry 43:5437–5444.

15. Zhao H, Butler E, Rodgers J, Spizzo T, Duesterhoeft S, Eide D (1998) Regulation of zinc homeostasis in yeast by binding of the ZAP1 transcriptional activator to zinc-responsive promoter elements. J. Biol. Chem. 273:28713–28720.

16. Wilson S, Bird AJ (2016) Zinc sensing and regulation in yeast model systems. Arch. Biochem. Biophys.

17. Wang Z, Feng LS, Matskevich V, Venkataraman K, Parasuram P, Laity JH (2006) Solution Structure of a Zap1 Zinc-responsive Domain Provides Insights into Metalloregulatory Transcriptional Repression in Saccharomyces cerevisiae. J. Mol. Biol. 357:1167–1183.

18. Qiao W, Mooney M, Bird AJ, Winge DR, Eide DJ (2006) Zinc binding to a regulatory zinc-sensing domain monitored in vivo by using FRET. Proc. Natl. Acad. Sci. 103:8674–8679.

19. Yao R, Li R, Huang Y (2023) Zinc homeostasis in Schizosaccharomyces pombe. Arch. Microbiol. 205:126.

20. Wilson S, Liu Y, Cardona-Soto C, Wadhwa V, Foster MP, Bird AJ (2019) The Loz1 transcription factor from Schizosaccharomyces pombe binds to Loz1 response elements and represses gene expression when zinc is in excess. Mol. Microbiol. 112:1701–1717.

21. Corkins ME, May M, Ehrensberger KM, Hu Y-M, Liu Y-H, Bloor SD, Jenkins B, Runge KW, Bird AJ (2013) Zinc finger protein Loz1 is required for zinc-responsive regulation of gene expression in fission yeast. Proc. Natl. Acad. Sci. U. S. A. 110:15371–6.

22. Ehrensberger KM, Corkins ME, Choi S, Bird AJ (2014) The double zinc finger domain and adjacent accessory domain from the transcription factor loss of zinc sensing 1 (Loz1) are necessary for DNA binding and zinc sensing. J. Biol. Chem. 289:18087–18096.

23. Auld DS (2001) Zinc coordination sphere in biochemical zinc sites. BioMetals.

24. Delaglio F, Grzesiek S, Vuister GeertenW, Zhu G, Pfeifer J, Bax A (1995) NMRPipe: A multidimensional spectral processing system based on UNIX pipes. J. Biomol. NMR 6.

25. Johnson BA From Raw Data to Protein Backbone Chemical Shifts Using NMRFx Processing and NMRViewJ Analysis. In: Methods in Molecular Biology.; 2018. pp. 257–310.

26. Shen Y, Bax A Protein Structural Information Derived from NMR Chemical Shift with the Neural Network Program TALOS-N. In: Methods in Molecular Biology.; 2015. pp. 17–32.

27. Palmer AG, Kroenke CD, Loria JP (2001) Nuclear magnetic resonance methods for quantifying microsecond-to-millisecond motions in biological macromolecules. Methods Enzymol. 339:204–38.

28. Hansen DF, Vallurupalli P, Kay LE (2008) An Improved 15 N Relaxation Dispersion Experiment for the Measurement of Millisecond Time-Scale Dynamics in Proteins †. J. Phys. Chem. B 112:5898–5904.

29. Kleckner IR, Foster MP (2012) GUARDD: user-friendly MATLAB software for rigorous analysis of CPMG RD NMR data. J. Biomol. NMR 52:11–22.

30. Waterhouse A, Bertoni M, Bienert S, Studer G, Tauriello G, Gumienny R, Heer FT, De Beer TAP, Rempfer C, Bordoli L, et al. (2018) SWISS-MODEL: homology modelling of protein structures and complexes. Nucleic Acids Res. 46.

31. Houbaviy HB, Usheva A, Shenk T, Burley SK (1996) Cocrystal structure of YY1 bound to the adeno-associated virus P5 initiator. Proc. Natl. Acad. Sci. U. S. A. 93:13577–13582.

32. Hudson NO, Whitby FG, Buck-Koehntop BA (2018) Structural insights into methylated DNA recognition by the C-terminal zinc fingers of the DNA reader protein ZBTB38. J. Biol. Chem. 293:19835–19843.

33. Li S, Olson WK, Lu X-J (2019) Web 3DNA 2.0 for the analysis, visualization, and modeling of 3D nucleic acid structures. Web Serv. Issue Publ. Online 47.

34. Stein A, Kortemme T (2013) Improvements to Robotics-Inspired Conformational Sampling in Rosetta Zhang Y, editor. PLoS ONE 8:e63090.

35. Touw WG, Van Beusekom B, Evers JMG, Vriend G, Joosten RP (2016) Validation and correction of ZnCysxHisy complexes. Acta Crystallogr. Sect. Struct. Biol. 72:1110–1118.

36. Van Zundert GCP, Rodrigues JPGLM, Trellet M, Schmitz C, Kastritis PL, Karaca E, Melquiond ASJ, Van Dijk M, De Vries SJ, Bonvin AMJJ (2016) The HADDOCK2.2 Web Server: User-Friendly Integrative Modeling of Biomolecular Complexes. J. Mol. Biol. 428:720–725.

37. Schubert M, Labudde D, Oschkinat H, Schmieder P (2002) A software tool for the prediction of Xaa-Pro peptide bond conformations in proteins based on 13C chemical shift statistics. J. Biomol. NMR 24:149–54.

38. Shen Y, Bax A (2010) Prediction of Xaa-Pro peptide bond conformation from sequence and chemical shifts. J. Biomol. NMR 46:199–204.

39. Alderson TR, Lee JH, Charlier C, Ying J, Bax A (2018) Propensity for cis -Proline Formation in Unfolded Proteins. ChemBioChem 19:37–42.

40. Persikov A V., Osada R, Singh M (2009) Predicting DNA recognition by Cys2His2 zinc finger proteins. Bioinformatics 25:22–29.

41. Farrow NA, Zhang O, Forman-Kay JD, Kay LE (1994) A heteronuclear correlation experiment for simultaneous determination of 15N longitudinal decay and chemical exchange rates of systems in slow equilibrium. J. Biomol. NMR 4:727–734.

42. Reimer U, Scherer G, Drewello M, Kruber S, Schutkowski M, Fischer G (1998) Side-chain effects on peptidyl-prolyl cis/trans isomerisation. J. Mol. Biol. 279:449–460.

43. Grathwohl C, Wüthrich K (1981) Nmr studies of the rates of proline cis - trans isomerization in oligopeptides. Biopolymers 20:2623–2633.

44. Baxter NJ, Williamson MP (1997) Temperature dependence of 1H chemical shifts in proteins. J. Biomol. NMR 9:359–69.

45. Trainor K, Palumbo JA, MacKenzie DWS, Meiering EM (2020) Temperature dependence of NMR chemical shifts: Tracking and statistical analysis. Protein Sci. 29:306–314.

46. Cordier F, Grzesiek S (2002) Temperature-dependence of protein hydrogen bond properties as studied by high-resolution NMR 1 1Edited by P. E. Wright. J. Mol. Biol. 317:739–752.

47. Rehm T, Huber R, Holak TA (2002) Application of NMR in Structural Proteomics. Structure 10:1613–1618.

48. Tjandra N, Feller SE, Pastor RW, Bax A (1995) Rotational diffusion anisotropy of human ubiquitin from 15N NMR relaxation. J. Am. Chem. Soc. 117:12562–12566.

49. Tjandra N, Garrett DS, Gronenborn AM, Bax A, Clore GM (1997) Defining long range order in NMR structure determination from the dependence of heteronuclear relaxation times on rotational diffusion anisotropy. Nat. Struct. Biol. 4:443–449.

50. Fushman D, Xu R, Cowburn D (1999) Direct Determination of Changes of Interdomain Orientation on Ligation: Use of the Orientational Dependence of 15 N NMR Relaxation in Abl SH(32) †. Biochemistry 38:10225–10230.

51. Bruschweiler R, Liao X, Wright P (1995) Long-range motional restrictions in a multidomain zinc-finger protein from anisotropic tumbling. Science 268:886–889.

52. Kleckner IR, Foster MP (2011) An introduction to NMR-based approaches for measuring protein dynamics. Biochim. Biophys. Acta - Proteins Proteomics 1814:942–968.

53. Foster MP, Wuttke DS, Clemens KR, Jahnke W, Radhakrishnan I, Tennant L, Reymond M, Chung J, Wright PE (1998) Chemical shift as a probe of molecular interfaces: NMR studies of DNA binding by the three amino-terminal zinc finger domains from transcription factor IIIA. J. Biomol. NMR 12:51–71.

54. Zhu R, Song Y, Liu H, Yang Y, Wang S, Yi C, Chen PR (2017) Allosteric histidine switch for regulation of intracellular zinc(II) fluctuation. Proc. Natl. Acad. Sci. 114:13661–13666.

55. Chandra BR, Yogavel M, Sharma A (2007) Structural Analysis of ABC-family Periplasmic Zinc Binding Protein Provides New Insights Into Mechanism of Ligand Uptake and Release. J. Mol. Biol.

56. Bonchuk AN, Georgiev PG (2024) C2H2 proteins: Evolutionary aspects of domain architecture and diversification. BioEssays 46:2400052.

57. Tsui V, Zhu L, Huang TH, Wright PE, Case DA (2000) Assessment of zinc finger orientations by residual dipolar coupling constants. J. Biomol. NMR 16:9–21.

58. Bédard M, Roy V, Montagne M, Lavigne P (2017) Structural Insights into c-Myc-interacting Zinc Finger Protein-1 (Miz-1) Delineate Domains Required for DNA Scanning and Sequence-specific Binding. J. Biol. Chem. 292:3323–3340.

59. Elrod-Erickson M, Pabo CO (1999) Binding studies with mutants of Zif268. Contribution of individual side chains to binding affinity and specificity in the Zif268 zinc finger-DNA complex. J. Biol. Chem. 274:19281–19285.

60. Peisach E, Pabo CO (2003) Constraints for Zinc Finger Linker Design as Inferred from X-ray Crystal Structure of Tandem Zif268–DNA Complexes. J. Mol. Biol. 330:1–7.

61. Gurung D, Danielson JA, Tasnim A, Zhang J-T, Zou Y, Liu J-Y (2023) Proline Isomerization: From the Chemistry and Biology to Therapeutic Opportunities. Biology 12:1008.

62. Boisvert O, Létourneau D, Delattre P, Tremblay C, Jolibois É, Montagne M, Lavigne P (2022) Zinc Fingers 10 and 11 of Miz-1 undergo conformational exchange to achieve specific DNA binding. Structure 30:623–636.e5.

63. Dovat S, Ronni T, Russell D, Ferrini R, Cobb BS, Smale ST (2002) A common mechanism for mitotic inactivation of C2H2 zinc finger DNA-binding domains. Genes Dev. 16:2985–2990.

64. Rizkallah R, Hurt MM (2009) Regulation of the transcription factor YY1 in mitosis through phosphorylation of its DNA-binding domain. Mol. Biol. Cell 20:4766–4776.

65. Rizkallah R, Alexander KE, Hurt MM (2011) Global mitotic phosphorylation of C2H2 zinc finger protein linker peptides. Cell Cycle 10:3327–3336.

66. Kluska K, Adamczyk J, Krężel A (2018) Metal binding properties, stability and reactivity of zinc fingers. Coord. Chem. Rev. 367:18–64.

67. Sebák F, Ecsédi P, Bermel W, Luy B, Nyitray L, Bodor A (2022) Selective 1Hα NMR Methods Reveal Functionally Relevant Proline cis/trans Isomers in Intrinsically Disordered Proteins: Characterization of Minor Forms, Effects of Phosphorylation, and Occurrence in Proteome. Angew. Chem. Int. Ed. 61:e202108361.

68. Belova E, Maksimenko O, Georgiev P, Bonchuk A (2022) The Essential Role of Prolines and Their Conformation in Allosteric Regulation of Kaiso Zinc Finger DNA-Binding Activity by the Adjacent C-Terminal Loop. Int. J. Mol. Sci. 23:15494.

69. Hanes SD (2015) Prolyl isomerases in gene transcription. Biochim. Biophys. Acta BBA - Gen. Subj. 1850:2017–2034.

70. Zhang X, Blumenthal RM, Cheng X (2024) Updated understanding of the protein–DNA recognition code used by C2H2 zinc finger proteins. Curr. Opin. Struct. Biol. 87:102836.

71. Bowers PM, Schaufler LE, Klevit RE (1999) A folding transition and novel zinc finger accessory domain in the transcription factor ADR1. Nat. Struct. Biol.

72. Kamashev DE, Balandina A V., Karpov VL (2000) Tramtrack protein-DNA interactions: A cross-linking study. J. Biol. Chem.

73. Jiang H, Daniels PJ, Andrews GK (2003) Putative zinc-sensing zinc fingers of metal-response element-binding transcription factor-1 stabilize a metal-dependent chromatin complex on the endogenous metallothionein-I promoter. J. Biol. Chem. 278:30394–30402.

74. Ogo OA, Tyson J, Cockell SJ, Howard A, Valentine RA, Ford D (2015) The Zinc Finger Protein ZNF658 Regulates the Transcription of Genes Involved in Zinc Homeostasis and Affects Ribosome Biogenesis through the Zinc Transcriptional Regulatory Element. Mol. Cell. Biol. 35:977–987.

75. Li Y, Kimura T, Laity JH, Andrews GK (2006) The Zinc-Sensing Mechanism of Mouse MTF-1 Involves Linker Peptides between the Zinc Fingers. Mol. Cell. Biol. 26:5580–5587.

76. Bernhardt ML, Kong BY, Kim AM, O’Halloran TV, Woodruff TK (2012) A Zinc-Dependent Mechanism Regulates Meiotic Progression in Mammalian Oocytes1. Biol. Reprod. 86:114, 1–10.

77. Maret W (2013) Zinc Biochemistry: From a Single Zinc Enzyme to a Key Element of Life. Adv. Nutr. 4:82–91.

78. Maret W (2017) Zinc in Cellular Regulation: The Nature and Significance of “Zinc Signals.” Int. J. Mol. Sci. 18:2285.

79. Rutherford KM, Lera-Ramírez M, Wood V (2024) PomBase: a Global Core Biodata Resource— growth, collaboration, and sustainability. Genetics 227:iyae007.

