## Supporting Information for "Conformational dynamics in specialized C_2_H_2_ zinc finger domains enable zinc-responsive gene repression in S. pombe"

### A rugged free energy landscape in specialized $C_2H_2$ zinc finger domains enables zinc-responsive gene repression in *S. pombe*

Vibhuti Wadhwa[1], Cameron Jamshidi[1], Kye Stachowski[1], Amanda J. Bird[2,*], Mark P. Foster[1,*]

[1]Department of Chemistry and Biochemistry, The Ohio State University, Columbus, Ohio.
[2]Department of Human Nutrition and Molecular Genetics, The Ohio State University, Columbus, Ohio.

Supporting Information

```
Loz1zf1:   468 YRCTECLQGFSRPSSLKIHTYSHT  491
               |.|..|.:.|..||:||:|...||
ZBTB38:      5 YACELCAKQFQSPSTLKMHMRCHT  30

Loz1zf2:   491 TGERPFVCDYAGCGKAFNVRSNMRRHQRIH  520
               |||:||.|.:.||||.|::..|:|.|.|||
YY1:        54 TGEKPFQCTFEGCGKRFSLDFNLRTHVRIH  83
```

*Figure S 1. Pairwise alignment of Loz1 zinc fingers against ZBTB38 (6e94) and YY1 (1ubd) structural zinc fingers, used as templates for homology modeling.*

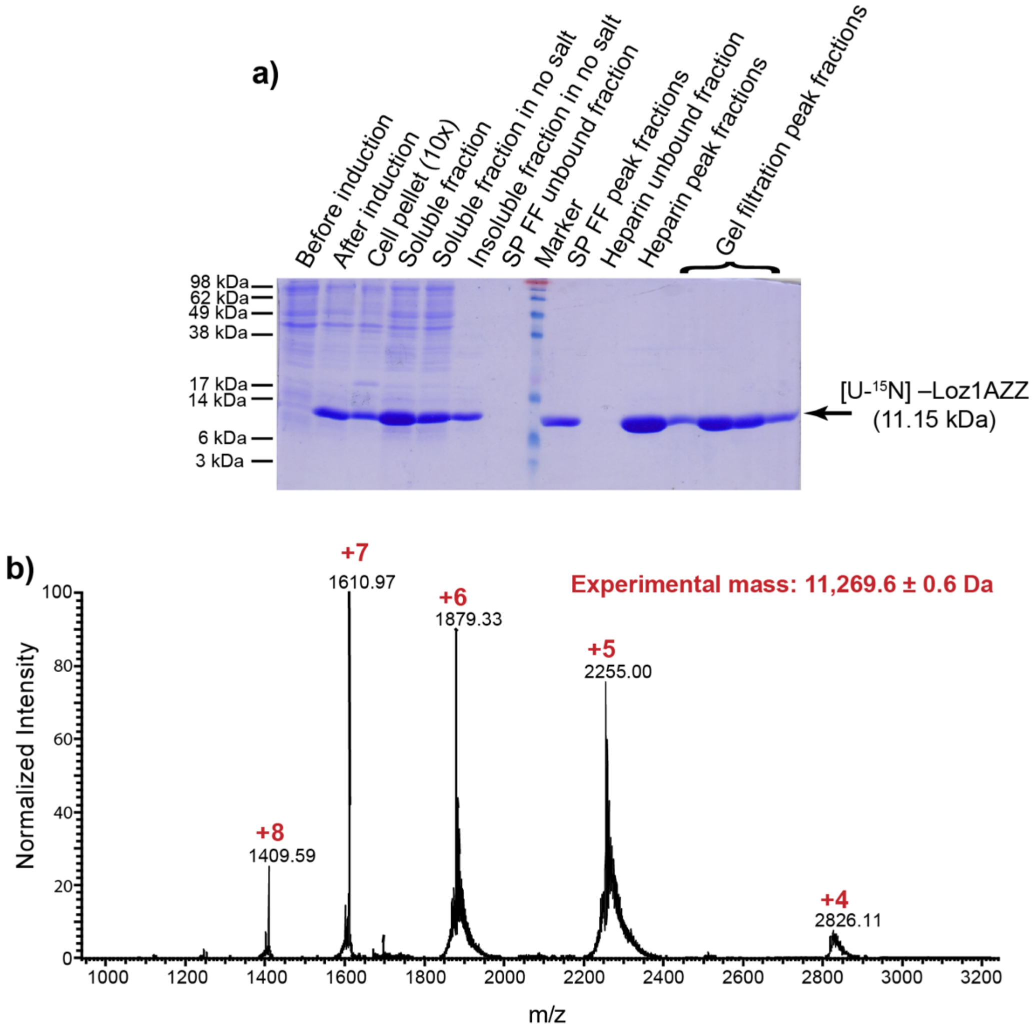


*Figure S 2.* ***Expression and purification of Loz1AZZ. a)*** *Samples at different stages of the purification of Loz1AZZ were subjected to electrophoresis on a 17% SDS gel and Coomassie-stained. The dashes and numbers to the left of the gel indicate the molecular masses of the marker.* ***b)*** *Native-MS spectrum of Loz1AZZ in 100 mM Ammonium Acetate showing charge state distribution and calculated molecular weight. Spectrum wes acquired on Q Exactive EMR Plus Orbitrap (Thermo) modified with a surface-induced dissociation (SID) device and a quadrupole. The expected molecular weight for [U-$^{15}$N]-Loz1AZZ is 11.28 kDa , when bound to two $Zn^{2+}$ ions.*

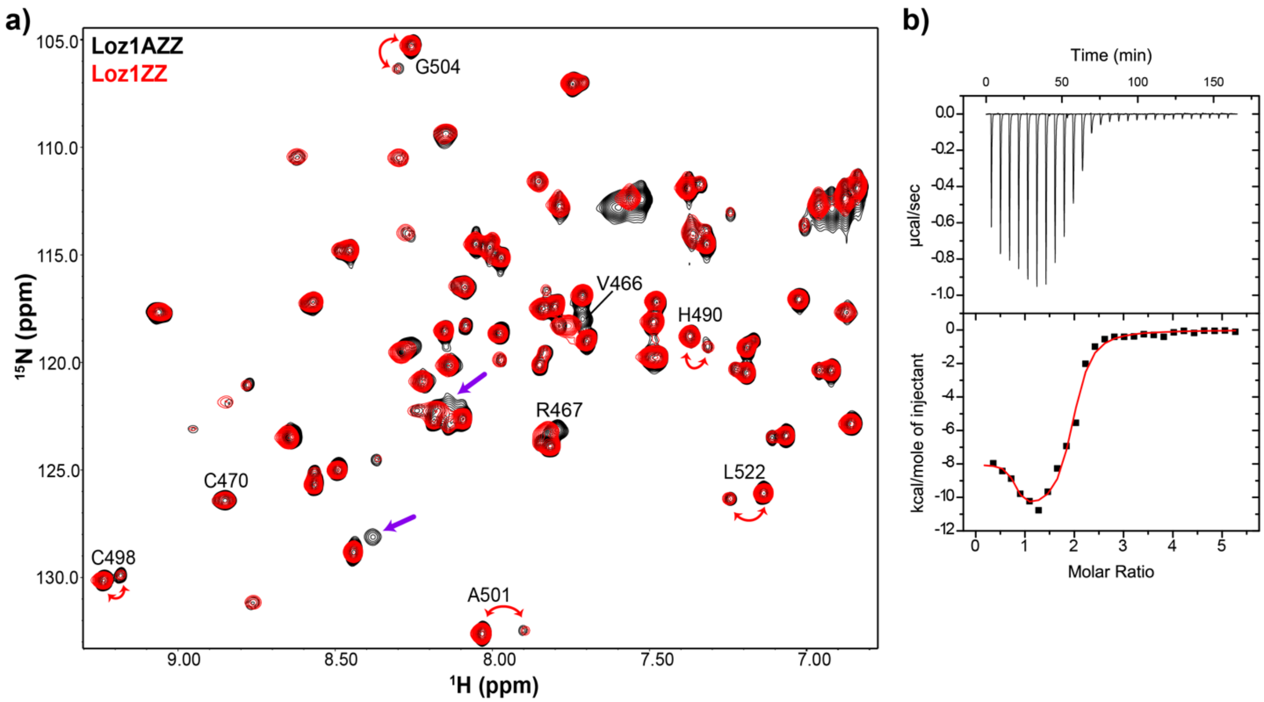


*Figure S 3. Supplementary figure 4: The accessory domain does not perturb Loz1 zinc finger structure and is not required for zinc-sensing. a) The {$^1$H}-$^{15}$N HSQC spectrum of Loz1ZZ (red) is mostly superimposable on that of Loz1AZZ (black), indicating that the zinc fingers fold independent of the accessory domain. Resonances attributed to the accessory domain are marked with an arrow. b) Representative ITC thermogram of zinc titration into apo-Loz1ZZ. The data were fit with a binding model with two independent sites. Best fit parameters are $n_1 = 0.73 \pm 0.07$, $K_{d1} = 3 \pm 4$ nM$^{-1}$, $\Delta H_1 = -8.02 \pm 0.42$ kcal mol$^{-1}$, $n_2 = 1.19 \pm 0.08$, $K_{d2} = 345 \pm 95$ nM, $\Delta H_2 = -10.98 \pm 0.45$ kcal mol$^{-1}$.*

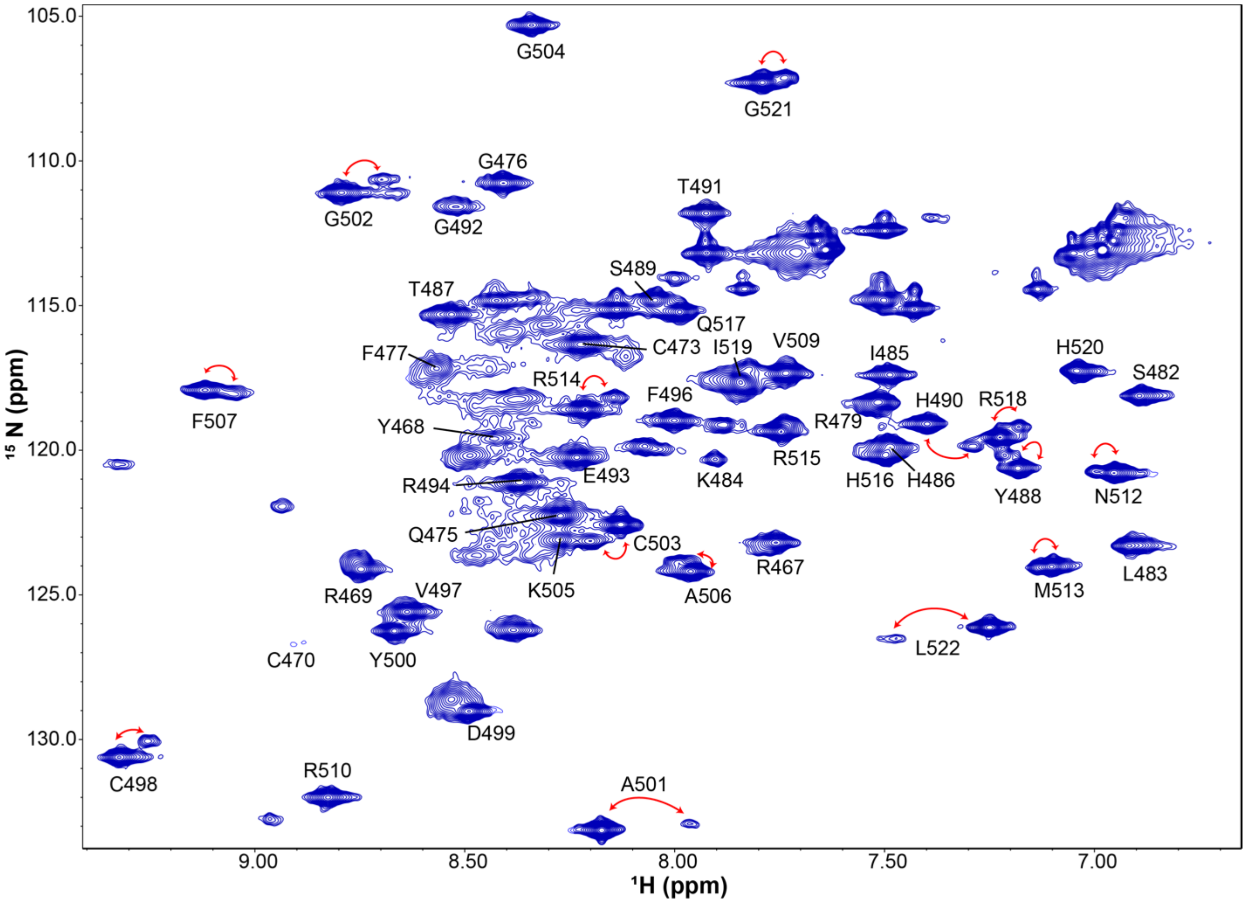


*Figure S 4. NMR spectra at low temperature reveal additional resonances attributable to an unstructured accessory domain. $^{1}H$ -$^{15}N$ HSQC of Loz1AZZ recorded at 273 K. Assigned signals from the zinc fingers are indicated and red arrows highlight resonances present in two slowly exchanging conformations. Low intensity broad signals, clustered between 8-8.5 $^{1}H$ ppm likely correspond to a poorly structured accessory domain.*

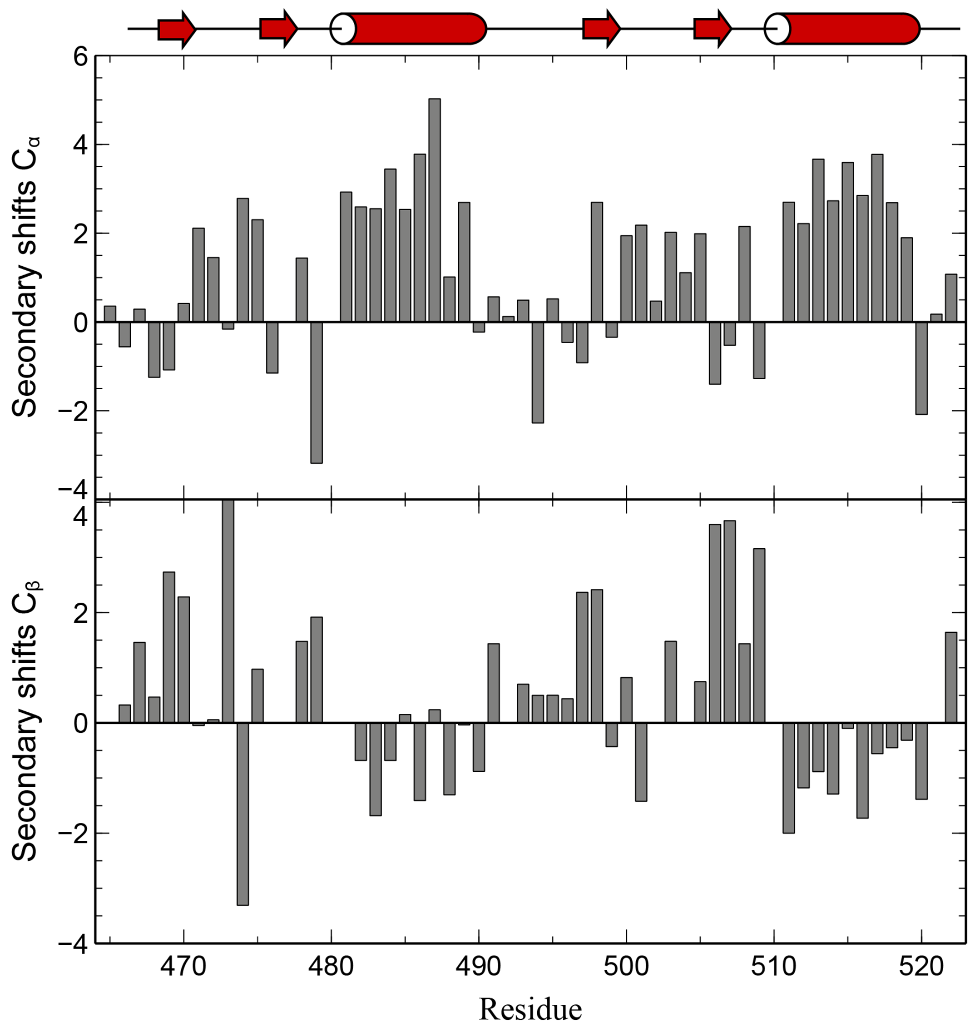


*Figure S 5.* ***Loz1 zinc finger chemical shifts are consistent with the canonical ββα fold.*** *Secondary $C^{\alpha}$ (top panel) and $C^{\beta}$ (bottom panel) chemical shifts as a function of residue number. Positive $C^{\alpha}$ secondary shits indicate a helical structure and negative values indicate strand. Behavior of $C^{\beta}$ atoms is opposite: negative values indicate helix and positive values are associated with strands. Schematic of predicted secondary structure fold is shown at the top.*

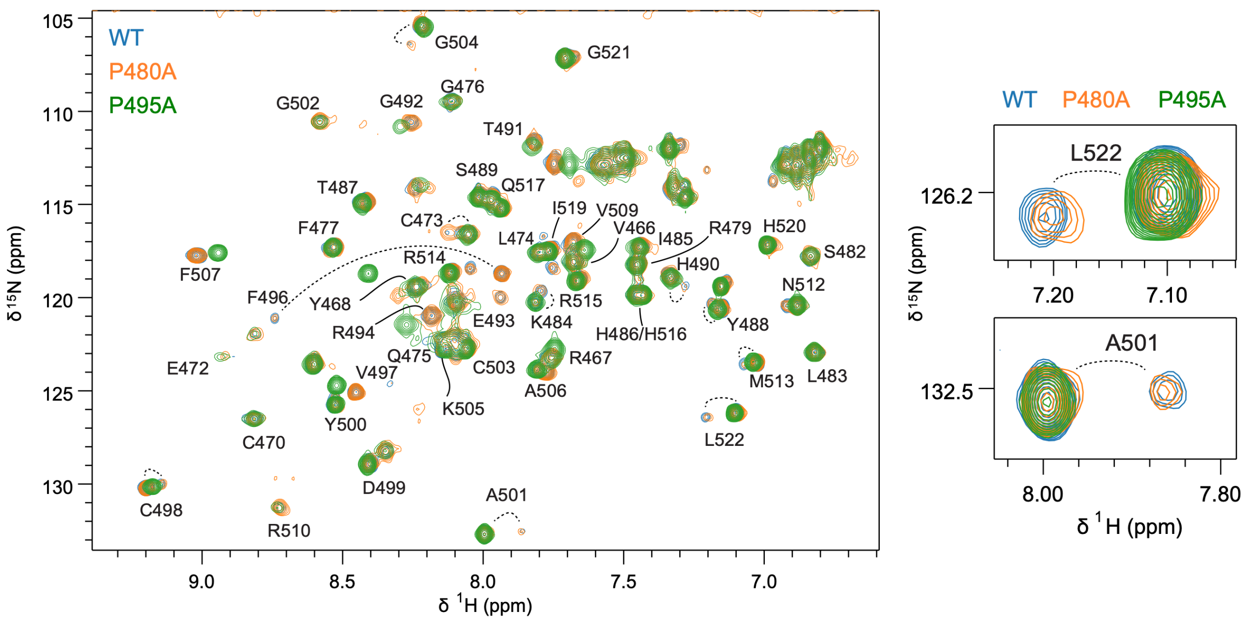


*Figure S 6. Slow cis-trans isomerization of Pro495 in the TGERP linker between ZF1 and ZF2 is responsible for signal doubling in the NMR spectra of Loz1AZZ. Overlaid $^{1}H$-$^{15}N$ 2D HSQC spectra of wild-type Loz1AZZ (blue), and variants with either Pro480 (orange) or Pro495 (green) mutated to Ala. Signal doubling observed in the WT spectrum (blue) persists in spectra of the P480A mutant (orange), but is absent in spectra of the P495A variant. Right, close-up view of signals from two residues diagnostic for the slow exchange behavior, Leu522 and Ala501. Spectra were recorded at 800 MHz, 25°C.*

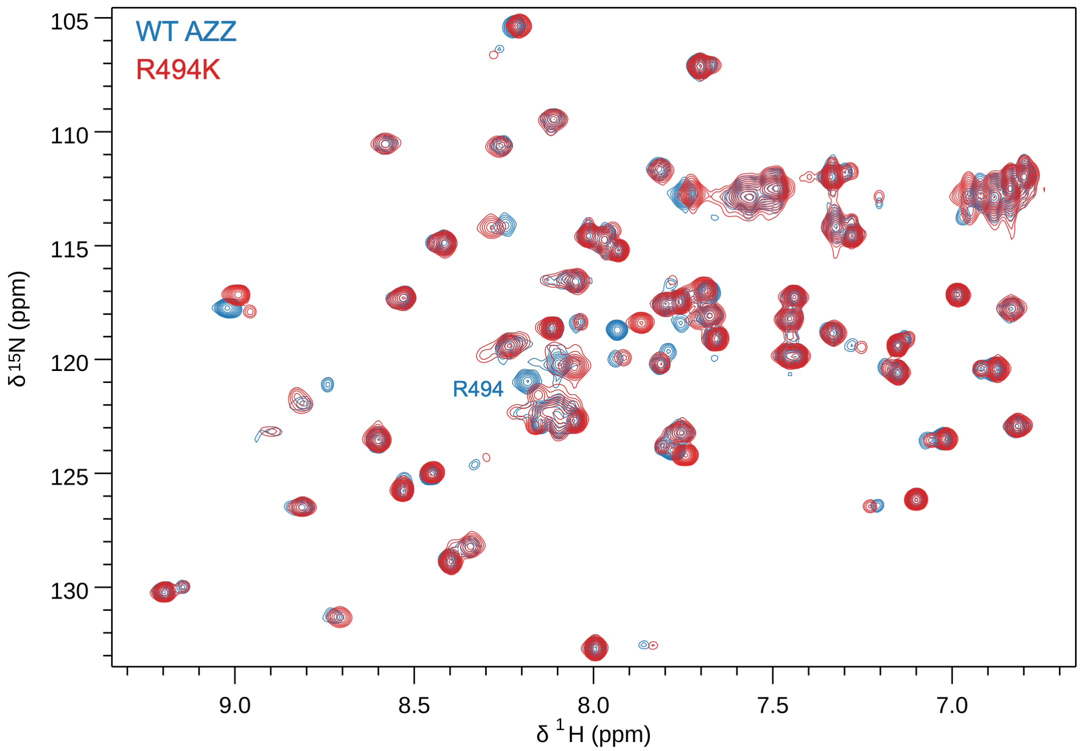


*Figure S 7. Replacement of Arg494 in the TGERP linker with Lys does not eliminate slow cis-trans X-Pro isomerization. Overlay of 2D HSQC spectra of WT (blue) and R494K-Loz1AZZ (red). Minor perturbations arising from the substitution are evident, but both spectra exhibit similar signal doubling, indicative of slow exchange between cis and trans peptide bond configurations.*

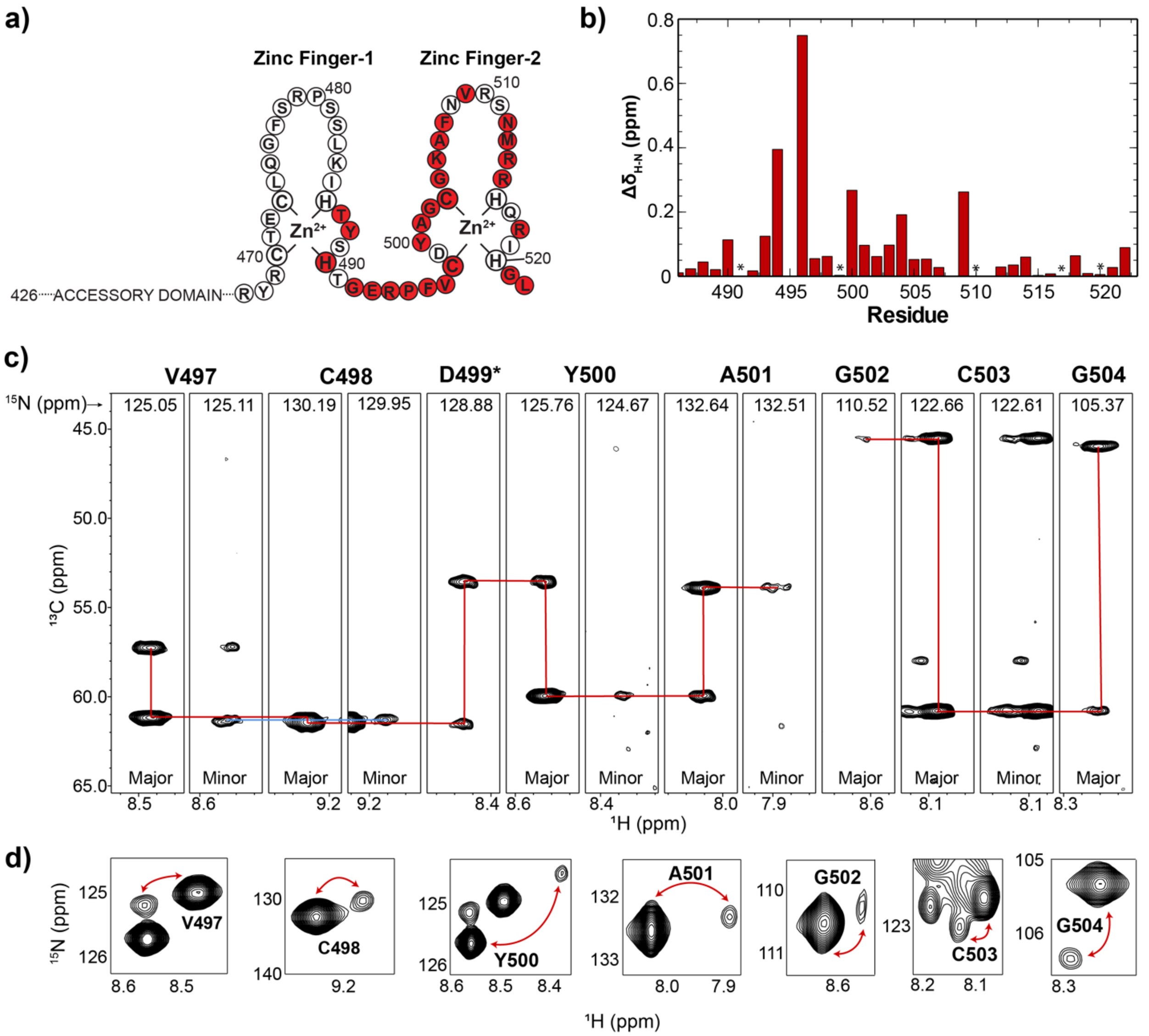


*Figure S 8.* ***Arg-Pro isomerization affects ZF1 and ZF2 residues. a)*** *Schematic of Loz1AZZ showing the zinc finger residues along with zinc coordination sites. Residues highlighted in red exhibit two signals from slowly exchanging conformations on the NMR time scale.* ***b)*** *Chemical shift perturbations plotted for residues highlighted in* ***a****. Assigned residues that do not display a second conformation are marked with *.* ***c)*** *Representative HNCA strips showing similarities and differences between Arg494-Pro495 trans and cis conformations of Loz1AZZ. A second set of signals were not observed for Asp499. Peaks for Gly502 minor species were below the noise level in the HNCA experiment.* ***d)*** *$^{1}H$-$^{15}N$ HSQC resonances of residues shown in* ***c****.*

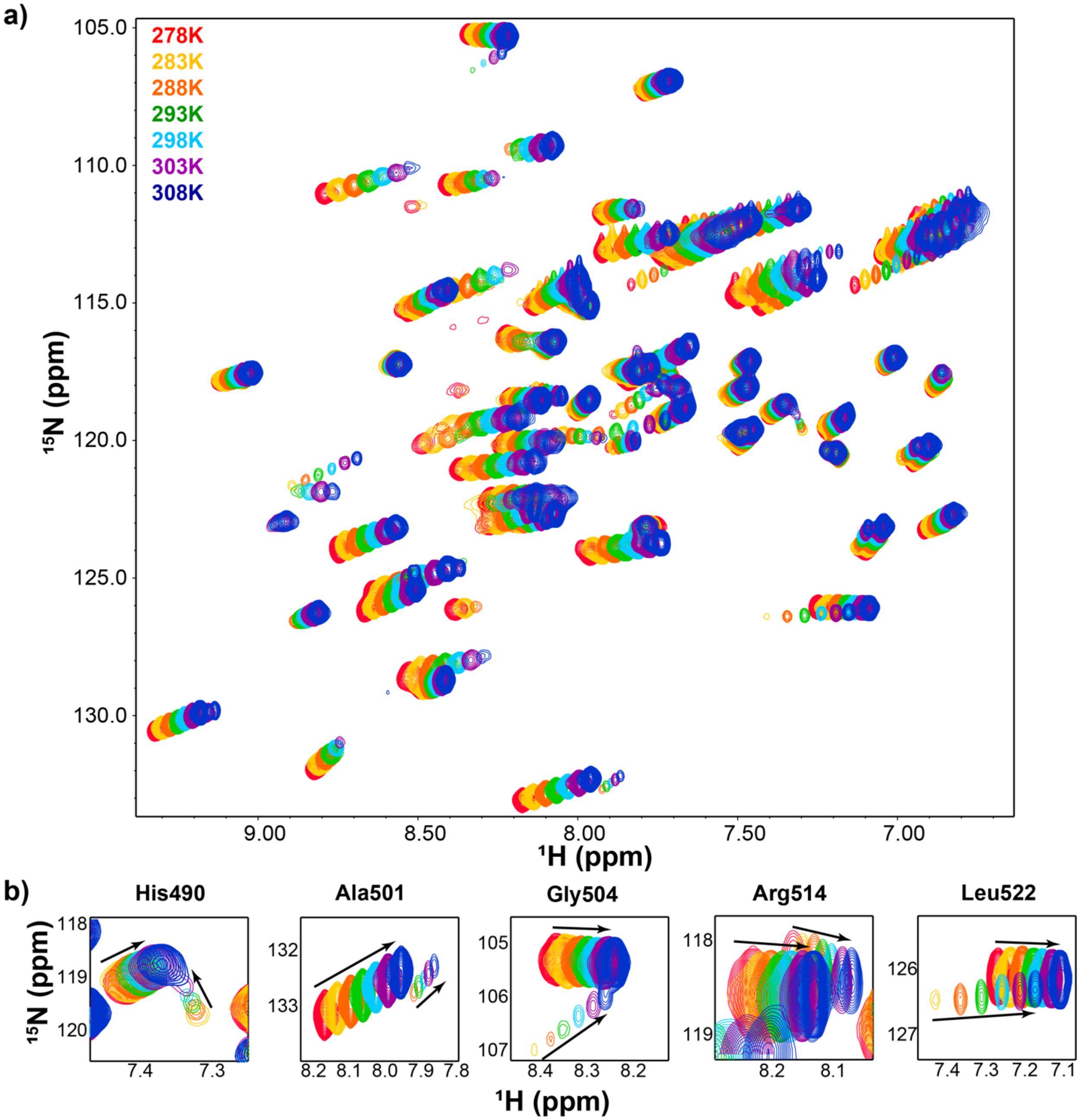


*Figure S 9.* ***Temperature sensitivity of Loz1AZZ amide signals. a)*** *Overlay of {$^{1}$H}-$^{15}$N spectra recorded over a range of temperatures for determining amide temperature coefficients.* ***b)*** *A subset of residues displaying peak pairs. Arrows show direction of chemical shit perturbations with increasing temperature.*

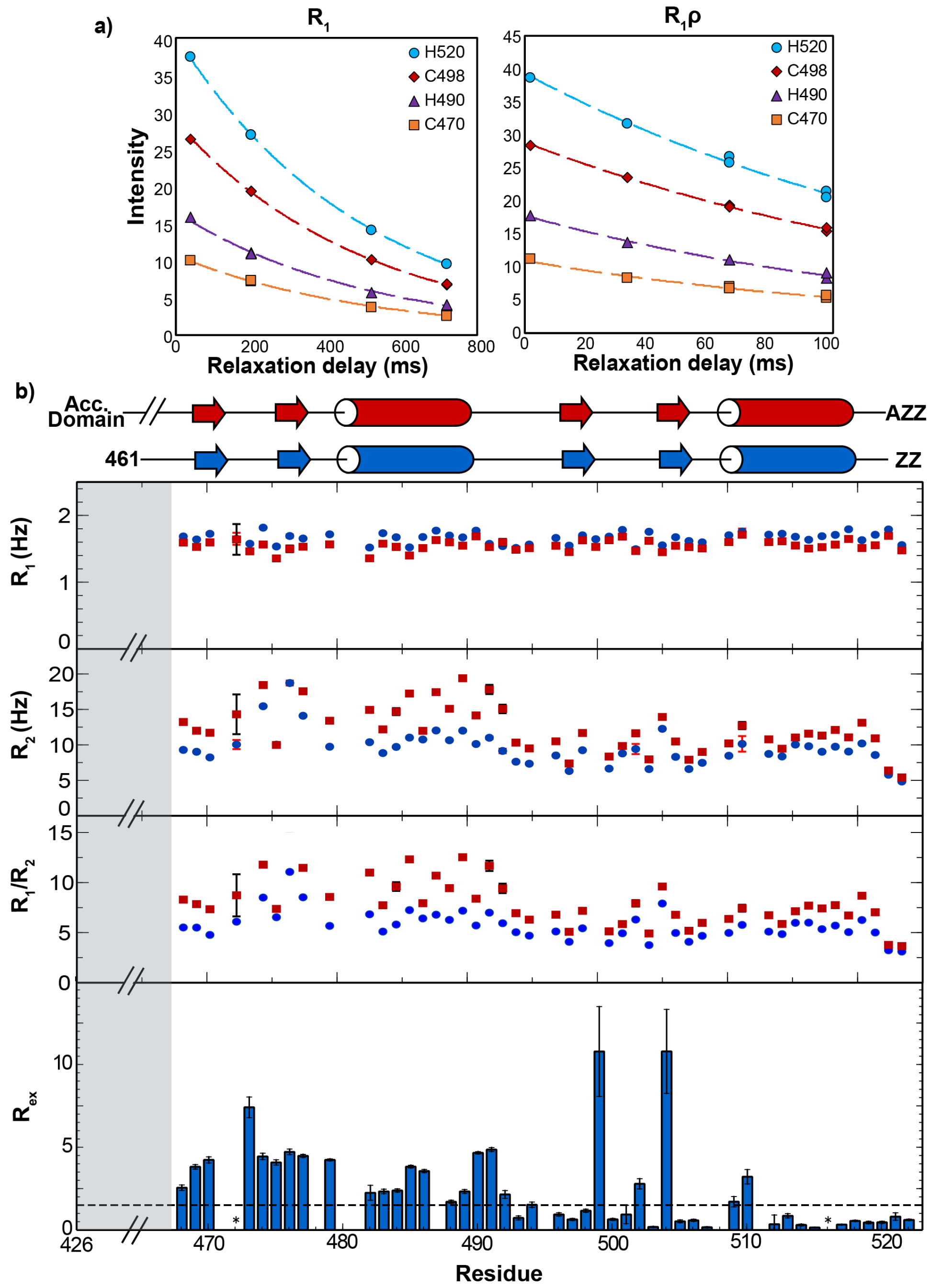


*Figure S 10.* ***Elevated $R_2$ relaxation is an intrinsic property of ZF1, independent of the accessory domain*****. *a)*** *$R_1$ and $R_1\rho$ representative relaxation decay curves for Loz1AZZ.* ***b)*** *Comparison of $R_1$, $R_2$ and $R_2/R_1$ values between Loz1AZZ (red squares) and Loz1ZZ (blue circles) zinc finger residues shows similar trends. Systematic offset is due to the difference in size of the protein constructs (11.16 kDa and 7.14 kDa for AZZ and ZZ, respectively). $R_1$ and $R_2$ errors were calculated from difference in intensities at relaxation delay duplicates. Bottom panel shows Loz1ZZ $R_{ex}$ values from 800MHz spectrometer data, obtained by fitting amide $^{15}N$ relaxation dispersion curves to a two-site exchange model. The (*) represents residues for which good fits of the dispersion curves could not be obtained or had Rex values close to zero.*

**Table S1:** Triple resonance experiments used for assigning amide and side-chain resonances of Loz1AZZ.

| **Experiment** | **SF** | | | **SW** | | | **SI** | | |
|---|---|---|---|---|---|---|---|---|---|
| Dimension | 1H | 15N | 13C | 1H | 15N | 13C | 1H | 15N | 13C |
| **HNCO** | 600.06 | 60.80 | 150.89 | 7500 | 1824.09 | 1660.03 | 2792 | 256 | 400 |
| **HNCA** | 600.06 | 60.80 | 150.89 | 7500 | 1824.09 | 3395.1 | 8192 | 200 | 376 |
| **HNCOCA** | 600.06 | 60.80 | 150.89 | 7500 | 1824.09 | 3395.1 | 2048 | 100 | 188 |
| **CBCACONH** | 600.06 | 60.80 | 150.89 | 7500 | 1824.09 | 8449.93 | 2048 | 232 | 392 |
| **HNCACB** | 600.06 | 60.80 | 150.89 | 7500 | 1824.09 | 8449.93 | 2048 | 232 | 408 |

| **Experiment** | **SF** | | | **SW** | | | **SI** | | | **Mixing time** |
|---|---|---|---|---|---|---|---|---|---|---|
| **Dimension** | 1H | 2nd | 3rd | 1H | 2nd | 3rd | 1H | 2nd | 3rd | |
| **(H)CCONH – TOCSY** | 800.13 | 81.09 (N) | 200.20 (C) | 7500 | 2433.09 | 12886.6 | 1024 | 120 | 256 | 14 ms; 10.4 kHz **$B_1$** |
| **H(CCO)NH – TOCSY** | 800.13 | 81.09 (N) | 800.13 (H) | 7500 | 2433.09 | 6402.05 | 8192 | 200 | 376 | 14 ms; 10.4 kHz **$B_1$** |
| **HCCH – TOCSY** | 800.13 | 201.20 (C) | 800.13 (H) | 10416.67 | 4426.41 | 5120.86 | 8192 | 248 | 624 | 15.6 ms; 10.4 kHz **$B_1$** |
| **HCCH – COSY** | 800.13 | 201.20 (C) | 800.13 (H) | 10416.67 | 4426.41 | 5120.86 | 8192 | 200 | 512 | |
| **15N – edited NOESY** | 850.28 | 86.17 (N) | 850.28 (H) | 11029.41 | 1723.37 | 11053.69 | 8192 | 160 | 840 | 200 ms |

SF: spectrometer frequency; SW, spectral width; SI; data size in time points.

**Table S2**. DNA oligonucleotide primers used to generate Loz1 mutants, P480A, P495A nad R494K.

P480A: F: CAAGGATTTTCTAGGGCTTCTAGTCTAAAAATTCATAC
P480A R: GTATGAATTTTTAGACTAGAAGCCCTAGAAAATCCTTG
P495A F: CCATACAGGAGAAAGGGCGTTTGTCTGCGATTAC
P495A R: GTAATCGCAGACAAACGCCCTTTCTCCTGTATGG
R494K F: CCCATACAGGAGAAAAGCCGTTTGTCTGCGATTAC
R494K R: GTAATCGCAGACAAACGGCTTTTCTCCTGTATGGG

F corresponds to the forward primer, R corresponds to the reverse primer.
mutagenic nucleotide(s) are underlined in the forward primers.
